# EASE: EM-Assisted Source Extraction from calcium imaging data

**DOI:** 10.1101/2020.03.25.007468

**Authors:** Pengcheng Zhou, Jacob Reimer, Ding Zhou, Amol Pasarkar, Ian Kinsella, Emmanouil Froudarakis, Dimitri V Yatsenko, Paul G Fahey, Agnes Bodor, JoAnn Buchanan, Dan Bumbarger, Gayathri Mahalingam, Russel Torres, Sven Dorkenwald, Dodam Ih, Kisuk Lee, Ran Lu, Thomas Macrina, Jingpeng Wu, Nuno da Costa, R. Clay Reid, Andreas S Tolias, Liam Paninski

## Abstract

Combining two-photon calcium imaging (2PCI) and electron microscopy (EM) provides arguably the most powerful current approach for connecting function to structure in neural circuits. Recent years have seen dramatic advances in obtaining and processing CI and EM data separately. In addition, several joint CI-EM datasets (with CI performed in vivo, followed by EM reconstruction of the same volume) have been collected. However, no automated analysis tools yet exist that can match each signal extracted from the CI data to a cell segment extracted from EM; previous efforts have been largely manual and focused on analyzing calcium activity in cell bodies, neglecting potentially rich functional information from axons and dendrites. There are two major roadblocks to solving this matching problem: first, dense EM reconstruction extracts orders of magnitude more segments than are visible in the corresponding CI field of view, and second, due to optical constraints and non-uniform brightness of the calcium indicator in each cell, direct matching of EM and CI spatial components is nontrivial.

In this work we develop a pipeline for fusing CI and densely-reconstructed EM data. We model the observed CI data using a constrained nonnegative matrix factorization (CNMF) framework, in which segments extracted from the EM reconstruction serve to initialize and constrain the spatial components of the matrix factorization. We develop an efficient iterative procedure for solving the resulting combined matching and matrix factorization problem and apply this procedure to joint CI-EM data from mouse visual cortex. The method recovers hundreds of dendritic components from the CI data, visible across multiple functional scans at different depths, matched with densely-reconstructed three-dimensional neural segments recovered from the EM volume. We publicly release the output of this analysis as a new gold standard dataset that can be used to score algorithms for demixing signals from 2PCI data. Finally, we show that this database can be exploited to (1) learn a mapping from 3d EM segmentations to predict the corresponding 2d spatial components estimated from CI data, and (2) train a neural network to denoise these estimated spatial components. This neural network denoiser is a stand-alone module that can be dropped in to enhance any existing 2PCI analysis pipeline.

## 1 Introduction

A fundamental goal of neuroscience is to understand the relationship between structure and function in neural circuits. Currently, arguably the most comprehensive available approach to link function with synaptic-resolution microanatomy is to perform two-photon calcium imaging (CI) followed by dense electron microscopy (EM) reconstruction of the same volume (Briggman et al., 2011; Bock et al., 2011; Lee et al., 2016; Vishwanathan et al., 2017; Hildebrand et al., 2017; Bae et al., 2018). This approach is highly labor intensive and expensive, and when successful provides highly scientifically valuable datasets.

Given the high value of this combined CI-EM data, we would like to extract as much information from these experiments as possible. Recent years have seen significant improvements in CI analysis (Mukamel et al., 2009; Pnevmatikakis et al., 2016; Pachitariu et al., 2016; Friedrich et al., 2017a,b; Petersen et al., 2018; Zhou et al., 2018; Buchanan et al., 2018; Soltanian-Zadeh et al., 2019; Giovannucci et al., 2019), and in parallel, major improvements in EM data acquisition and analysis (Helmstaedter et al., 2013; Kim et al., 2014; Kasthuri et al., 2015; Hayworth et al., 2015; Morgan et al., 2016; Ding et al., 2016; Zheng et al., 2017; Takemura et al., 2017; Januszewski et al., 2018; Motta et al., 2019).

However, automated analysis tools for joint CI-EM data are less mature; previous efforts have been largely manual and focused on analyzing calcium activity in cell bodies, neglecting potentially rich functional information from axons and dendrites. The goal of this paper is to develop analysis tools that provide a *dense fusion* of this joint CI-EM data, by using the densely-reconstructed EM data to help extract a more complete estimate of spatiotemporal functional signals from the CI data; see Figure 1 for a schematic illustration. Specifically, following (Pnevmatikakis et al., 2016) (and many of the other CI analysis papers referenced above), we want to decompose the observed calcium movie *Y* into the form

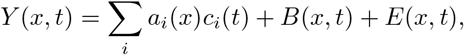

where *Y*(*x,t*) denotes the movie data at the *t*-th frame in the *x*-th pixel, *B*(*x,t*) is “background” signal (capturing highly-correlated activity that can not be separated into individual neuronal contributions), *E*(*x,t*) is temporally uncorrelated noise, *i* indexes neurons visible within the field of view (FOV), *c_i_*(*t*) is the calcium trace estimated from the *i*-th neuron, and *a_i_*(*x*) is the shape of the *i*-th neuron. The novel step here is that we add an additional constraint: each functional spatial component *a_i_* must be matched to a specific neuronal segment extracted from the EM reconstruction. If we are able to perform this matching correctly, then the EM segments serve as ground-truth, high spatial resolution, three-dimensional constraints on the shapes *a_i_*; these constraints should lead to better estimates of each *a_i_* and in turn the corresponding activity traces *c_i_*.

**Figure 1:**
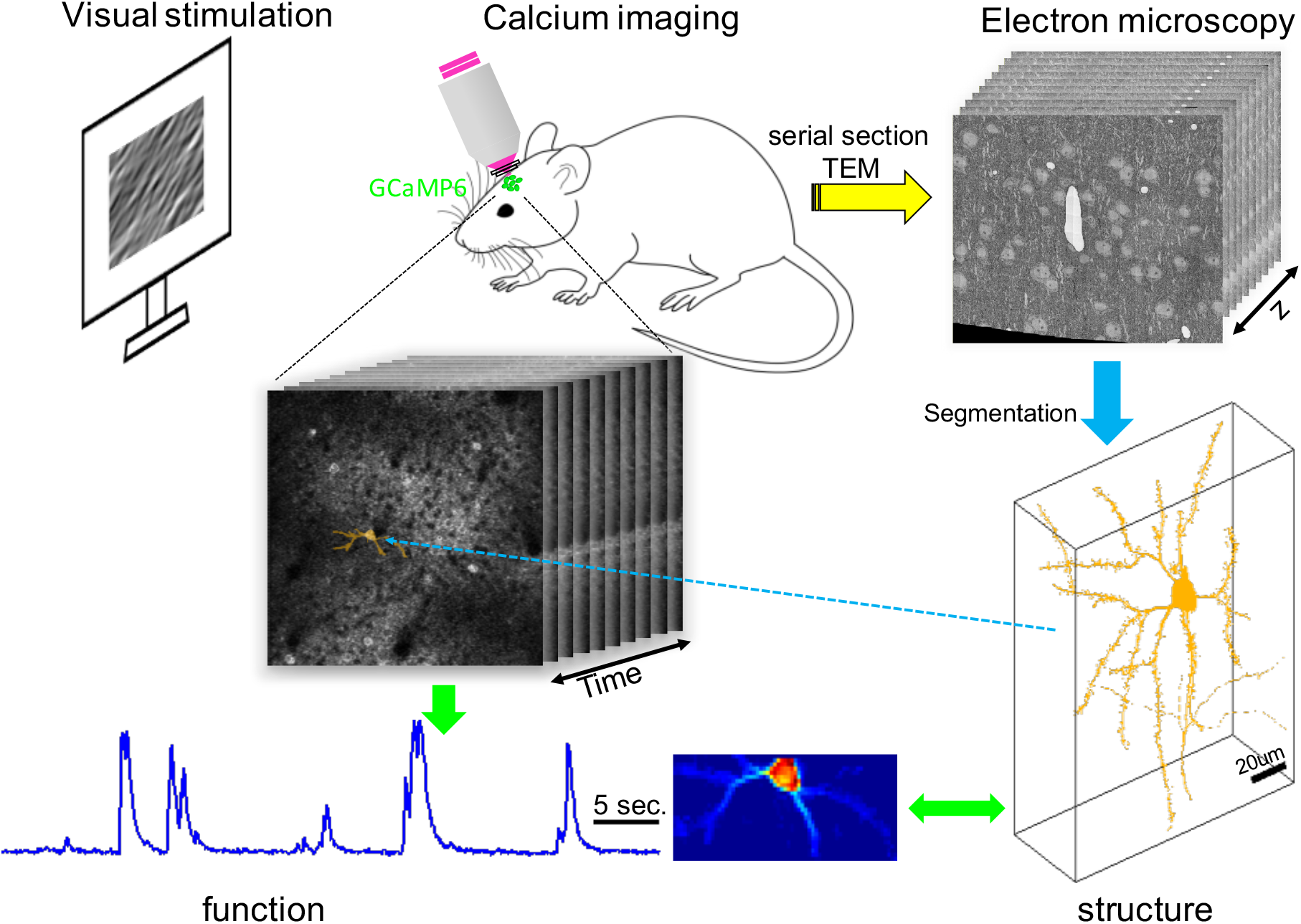
Schematic diagram of the experimental setup and overall analysis goal. *In vivo* calcium imaging and serial EM imaging were performed sequentially on the same brain volume in a single mouse. Segmenting the EM data provides the three-dimensional spatial structure of individual neurons at synaptic resolution. EASE combines these segments with the calcium imaging video to extract neural activity at single-neuron resolution, with each neuron visible in the functional calcium imaging movie matched with a corresponding neuron in the anatomical EM stack.

This matching problem is challenging for several reasons. First, the number of EM components in the FOV is large: in the data set we examine here, there are > 40000 EM segments intersecting a single FOV, but only a couple hundred good calcium components visible in the same FOV. Thus we have to solve a very sparse model selection problem — how to choose a couple hundred EM components that match the bright neurons in the calcium movie, then use these EM shapes to enforce constraints on the spatial components *a_i_* we extract from the CI movie. This would be at least conceptually easy if the spatial calcium components matched the EM components exactly; unfortunately, the second major problem is that the CI components *a_i_* typically represent only spatial subsets of the EM components, since the calcium indicator does not have a uniform brightness throughout the EM component. Finally, due to the large number of EM segments, overfitting is a potential problem: there may be many combinations of subsets of the EM components that can be added together to explain the observed data *Y*.

We note that there is a significant and growing literature on correlative light-electron microscopy (CLEM) (Maco et al., 2013, 2014a,b; de Boer et al., 2015; Blazquez-Llorca et al., 2015; Begemann and Galic, 2016; Lees et al., 2017; Drawitsch et al., 2018; Hoffman et al., 2020), but there are significant differences between the (functional, dynamic) CI data we analyze here and the (anatomical, static) light microscopy images handled in the CLEM literature. CI is used to simultaneously record population neurons’ activity at a fast temporal resolution, while in the context of CLEM the goal is to image fine static spatial structure using much longer imaging exposure times. Hence, CI data samples neurons’ spatial structures at lower spatial resolution and SNR per frame; on the other hand, by integrating time varying information over many imaging frames, CI offers the opportunity to demix overlapping neural shapes much more effectively than would be possible with a single fluorescent image, even one with high spatial resolution and SNR. Thus, in short, the CI-EM fusion problem we tackle here is distinct from the problems addressed so far in the CLEM literature, with its own unique challenges and opportunities.

We address these challenges by solving the matching and matrix factorization problems simultaneously. The basic approach is to start with the clearest matches (i.e., the EM segments that are most clearly visible in the CI data *Y*), then update the matrix factorization under the constraint that each CI component *a_i_* is contained within its matching EM segment, and then iterate this procedure, adding more matching components until no further good matches are available. We call the resulting algorithm EASE, for EM-Assisted Source Extraction.

In practice, EASE is scalable and effective: in a CI-EM dataset from mouse visual cortex EASE extracts the large majority of visible neuronal components from *Y* and automatically matches these components with the corresponding EM segments. Compared to the results without EM constraints, EASE leads to improved demixing of spatially-overlapping, correlated neurons and better recovery of small, low-SNR components. Finally, because the EM components are reconstructed in three dimensions, we can link the activity of extended dendritic segments that were functionally imaged across several different z depths at different times, thus providing multiple distinct views of the functional correlations within the imaged volume (Soudry et al., 2015).

The output of EASE is the factorization (*a_i_, c_i_, B*, and *E*) along with the matchings of each *a_i_* with the corresponding three-dimensional, high-resolution EM segment. We release these outputs publicly at this **site**. We believe this new annotated public dataset represents a valuable new “gold standard” that can in turn be used to score pipelines for demixing two-photon CI data. As an illustrative application, we show that this database can be exploited to (1) learn a mapping from 3d EM segmentations to predict the corresponding 2d spatial components estimated from CI data, and (2) train a neural network to denoise these estimated spatial components. This neural network denoiser is a stand-alone module that can be dropped in to enhance any existing two-photon CI analysis pipeline.

## 2 Results

### 2.1 Visualization and quantification of the extracted EM footprints

The first step of the EASE pipeline (described in full detail in the Methods section) is to project the three-dimensional high-resolution EM segments into the functional imaging planes. We begin in Figure 2 by examining the resulting EM “footprints” {*p_i_*} from a single scan. These footprints correspond to blurred and downsampled versions of the intersection of the full three-dimensional EM segments with the imaging plane: i.e., *p_i_* is a model of what cell *i* would look like in a two-photon scan if the brightness of the calcium indicator within the cell is uniform (see section 4.4 for details).

**Figure 2:**
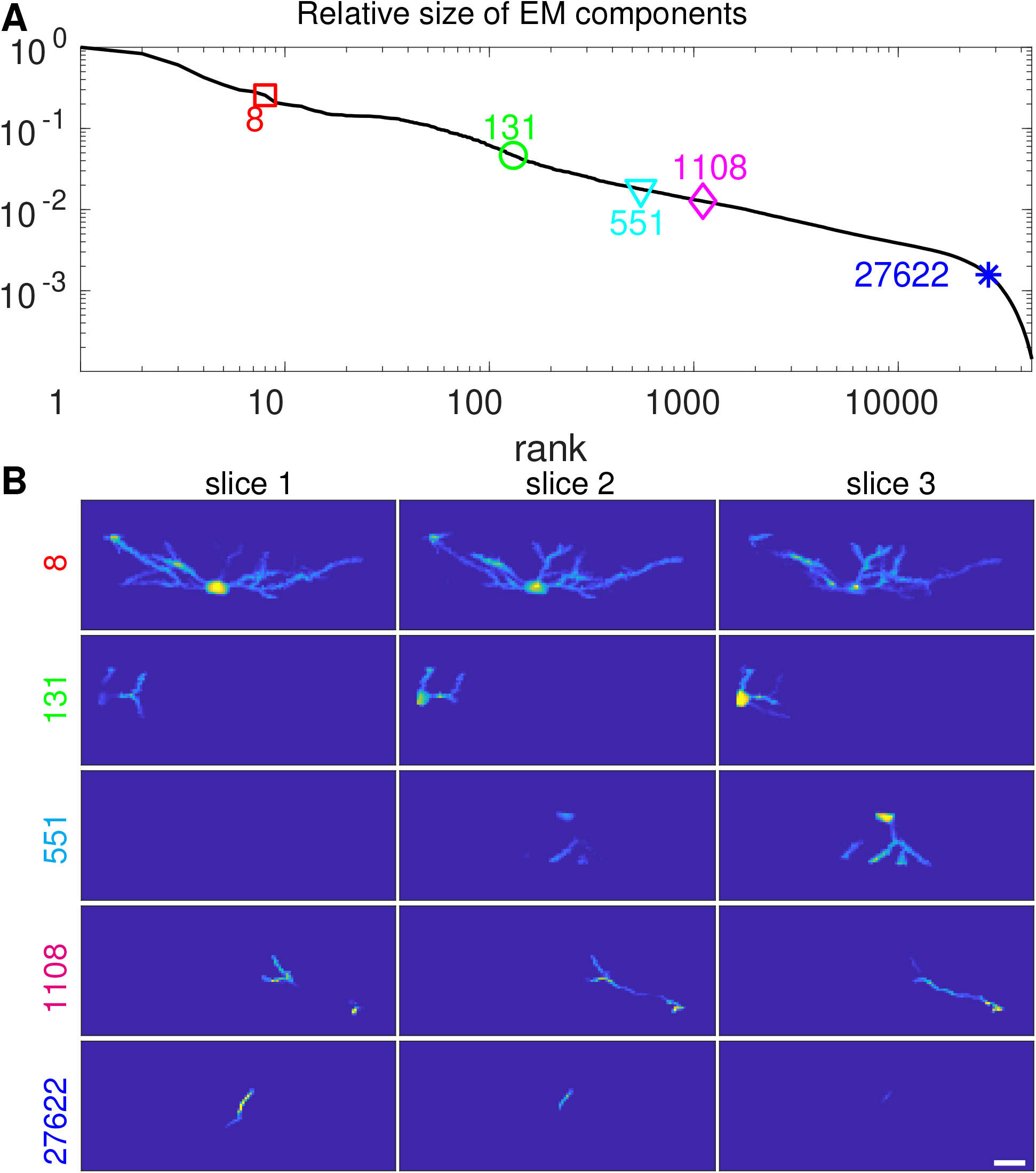
Example EM footprints *p_i_* from a single functional scan (recall that three imaging planes are obtained per functional scan in this dataset). (**A**) The relative size (measured as the *ℓ*_1_ norm, ∑_*x*_ *p_i_*(*x*)) of the EM footprints from this scan, in decreasing order. 5 example neurons were selected and their ranks were labeled with the same color. (**B**) The spatial footprints of the 5 selected examples. Note the wide ranges of sizes of these components; even the smallest component shown here (component 27622) could still plausibly contribute visible signal to the CI data *Y*. (Scale bar: 20 um).

A few points are immediate. First, there are many viable EM components visible in this scan: without any further information about cell type or calcium expression levels, there are tens of thousands of EM components with footprints *p_i_* which could plausibly contribute signal to the CI movie *Y*. Second, there is a wide range of footprint sizes in this dataset: when we sum over all pixels *x*,∑*_x_p_i_*(*x*) varies over more than two orders magnitude from the biggest *p_i_* to the smallest shown in Figure 2B. Finally, it is clear that non-somatic processes (e.g. dendrites) contribute many pixels to *p_i_* — in fact the large majority of pixels in the cell segments shown in Figure 2B are non-somatic, and therefore we can expect much of the signal in the CI data *Y* to be non-somatic as well.

### 2.2 Extraction of neural components from a single scan

Next we applied the EASE pipeline (described in algorithm (1) of the Methods section) to process data from a single scan (video size 58 × 129 × 3 *voxels* × 8900 *frames).* The details of all iterations are summarized in Table S1 (see Supplementary Information 5.1). The automated portion of the pipeline required ~ 6 minutes on a desktop computer. The pipeline yielded 173 components, among which an estimated 25 components included cell bodies (identified by eye), while the remaining 148 components included only non-somatic processes (likely dendrites). Figure 3AB shows 20 example components, evenly drawn from the top 100 components (ordered by brightness). The inferred spatial CI footprints *a_i_* resemble the corresponding EM footprints *p_i*_*, as desired. No large remaining signals were apparent upon visual examination of the residual video (S1 Video) or the peak-to-noise (PNR) image of the residual (Figure 3C).

**Figure 3:**
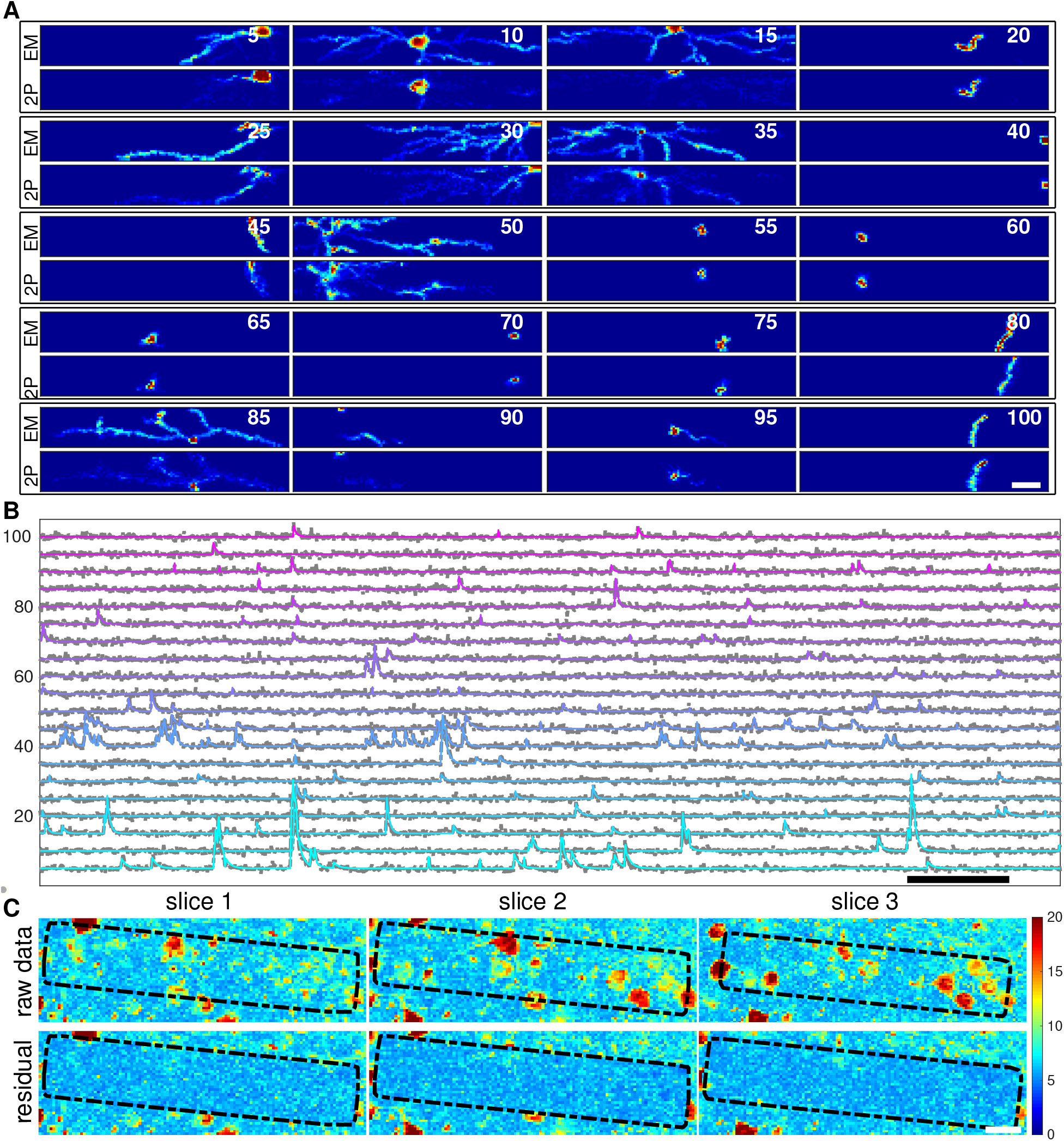
EASE results on a single CI scan. (**A**) spatial footprints of example neurons (rank in brightness order indicated in each subpanel). Scale bar: 20 um. Note that the EM footprints *p_i_* and the estimated CI components *a_i_* match each other well. (**B**) temporal traces of the example neurons shown in A. Color traces are denoised activity; gray traces are raw. All traces were normalized to have the same noise level. Scale bar: 10 seconds. (**C**) The peak-to-noise ratio (PNR) images before and after extracting neurons within the EM volume. Black rectangle indicates the boundaries of the EM volume; note that no bright signals remain in the residual within this region.

In Figure 4 we examine all of the estimated spatial components *a_i_* together. The density of the recovered neurons is quite high, so to improve the clarity of this visualization we broke the components into five groups, sorted by the confidence scores defined in section 4.9. We conclude that somatic components tend to be identified with the highest confidence; then processes that branch within the field of view (leading to components *a_i_* with a large total number of pixels); then finally, processes (including many apical dendrites) that cut through the imaging plane so that only a few pixels are visible lead to the lowest relative confidence scores. For these components (“other,” bottom row, Figure 4) we choose not to assign a definite match, to avoid corrupting downstream analyses with an overly confident but mistaken match.

**Figure 4:**
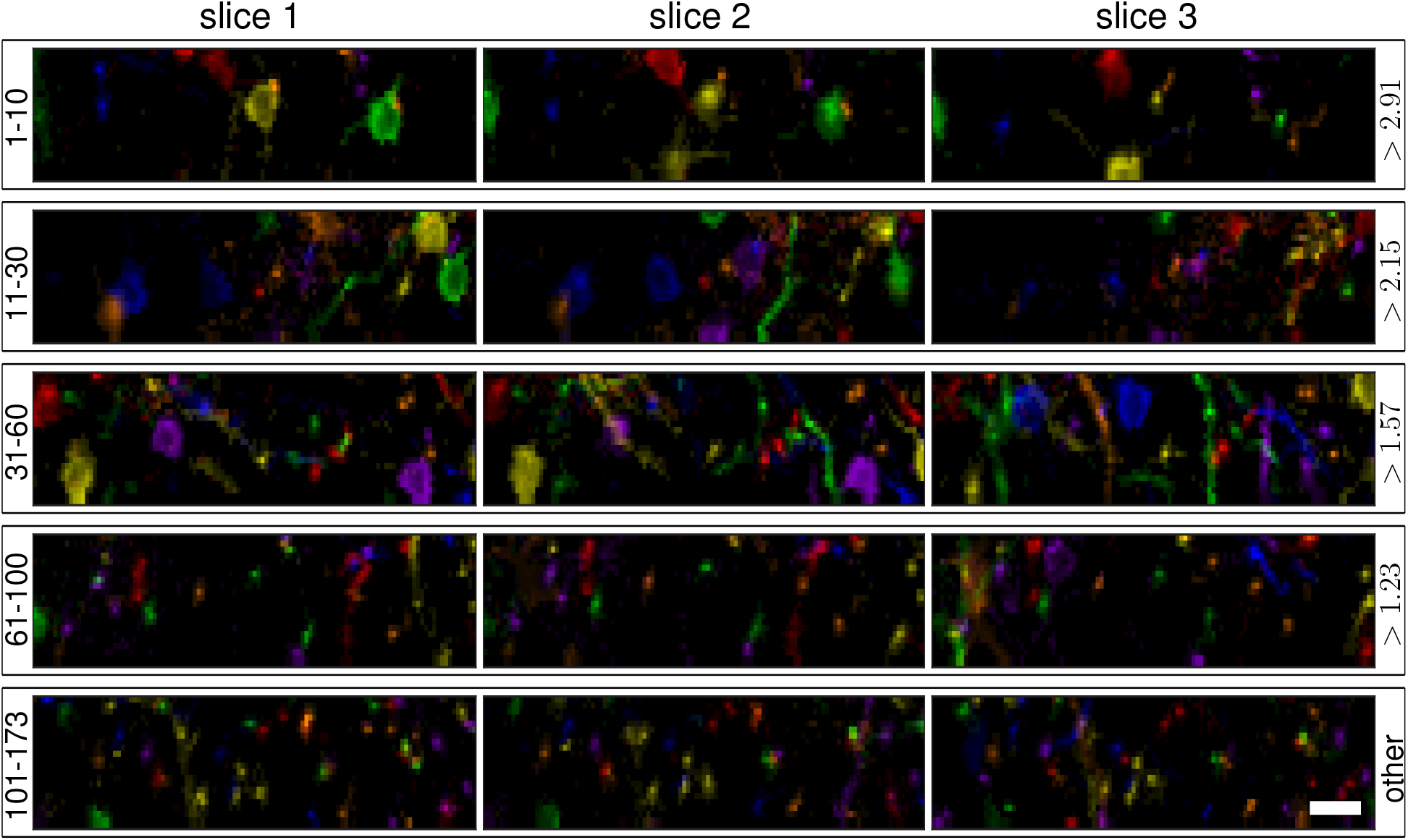
Spatial components extracted from a single scan. Each component is indicated with a single color; the estimated *a_i_*(x) scales the intensity of this color at each pixel x. Components were ordered according to their confidence scores (defined in section 4.9); each row here shows a subset of the spatial components *a_i_*, divided into groups according to their confidence rank (confidence ranks shown on the left; confidence score range shown on the right). Note that confidence scores for somatic components (top rows) tend to be higher than scores for small apical dendritic components, which dominate the bottom rows here. Scale bar: 20 um.

The EM information exploited by EASE provides a strong prior on the shapes of the targeted neurons, enabling the algorithm to demix neurons with strong spatial overlaps. This point is illustrated in Figure 5. We performed a simple clustering to order neurons with strong spatial overlaps next to each other, leading to a strong blockwise structure in the matrix formed by computing correlations between each spatial component *a_i_* (Figure 5A). Importantly, the corresponding temporal correlations (Figure 5B) did not display the same strong blockwise structure — i.e., components with high spatial overlap did not necessarily display correspondingly strong temporal correlations (which might have indicated problems demixing spatially overlapping signals in *Y*). Figure 5CD examines a block of four components with high spatial overlap: again, no problematic demixing issues are visible in either the spatial or temporal components recovered here. (See S2 Video for a zoomed-in depiction.)

**Figure 5:**
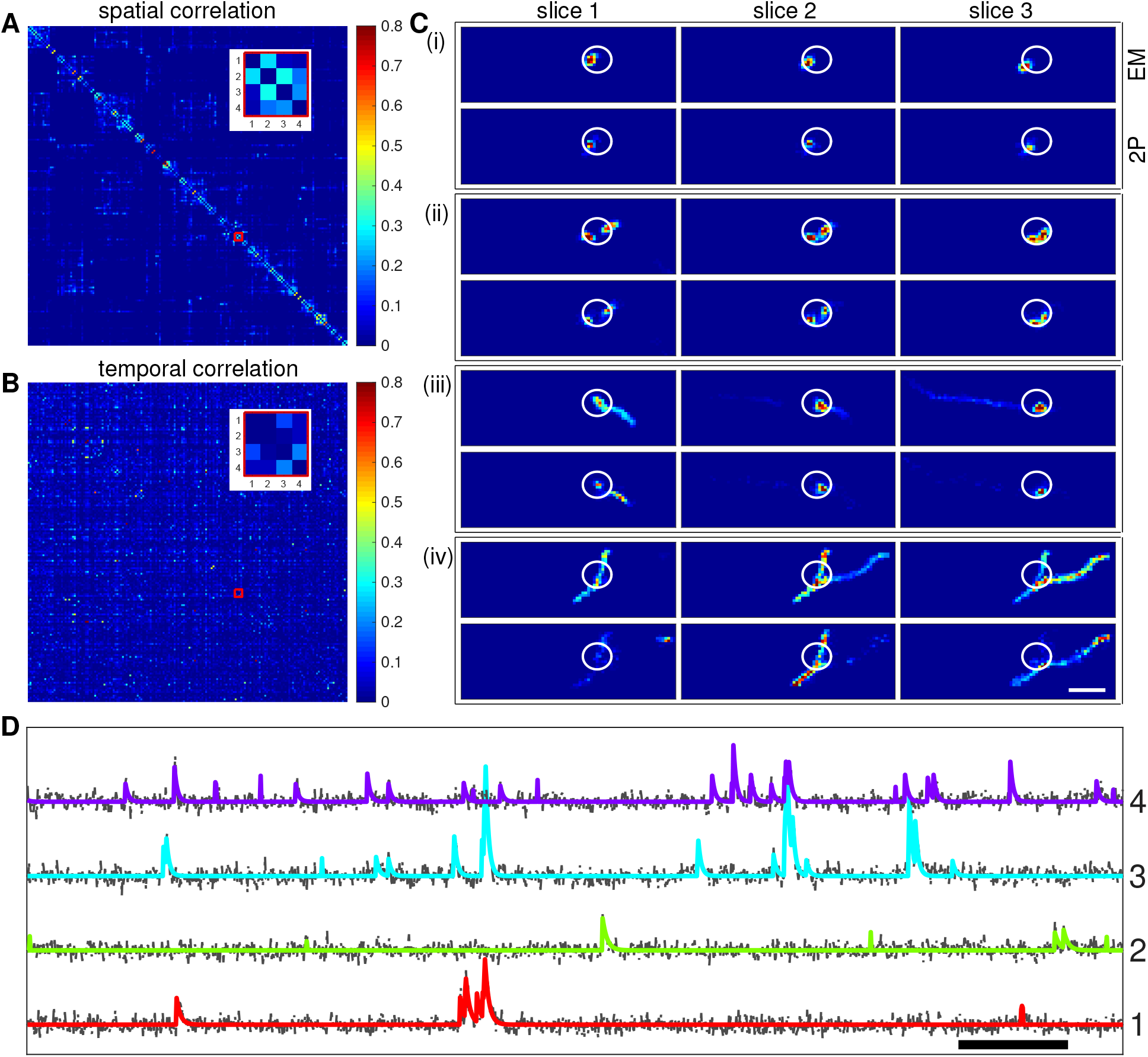
EASE can demix spatially overlapped neurons’ signals. (**A**) pairwise spatial correlation between all extracted components (auto-correlations excluded to improve visibility). We used hierarchical clustering of the spatial components to place spatially overlapping neurons together. Note the large degree of overlap visible (c.f. Figure 4). (**B**) pairwise temporal correlation, using the same ordering of components as in A. (**C**) spatial footprints of the selected 4 example neurons shown in the inset in panels A and B. Scale bar: 20 um. White circles simply indicate the same location in each panel, to aid comparisons. (**D**) temporal traces of the selected neurons; conventions as in Figure 3B. Scale bar: 10 seconds.

Conversely, we noticed some extracted components with highly correlated temporal traces but completely disconnected spatial components (Figure 6). By overlaying these extracted footprints onto the correlation image between the raw video *Y* and the average of the temporal traces *c_i_*, we found that these components actually correspond to different segments of the same neuron (Figure 6A), although they were not connected within the EM volume. These cases were rare and typically occurred at the boundary of the EM volume, where multiple dendrites from the neuron entered the EM volume at different (spatially disconnected) locations. It would not be feasible to re-join these components with purely anatomical methods using the EM data, but the functional information from the CI data makes it possible to rejoin these severed connections with high confidence.

**Figure 6:**
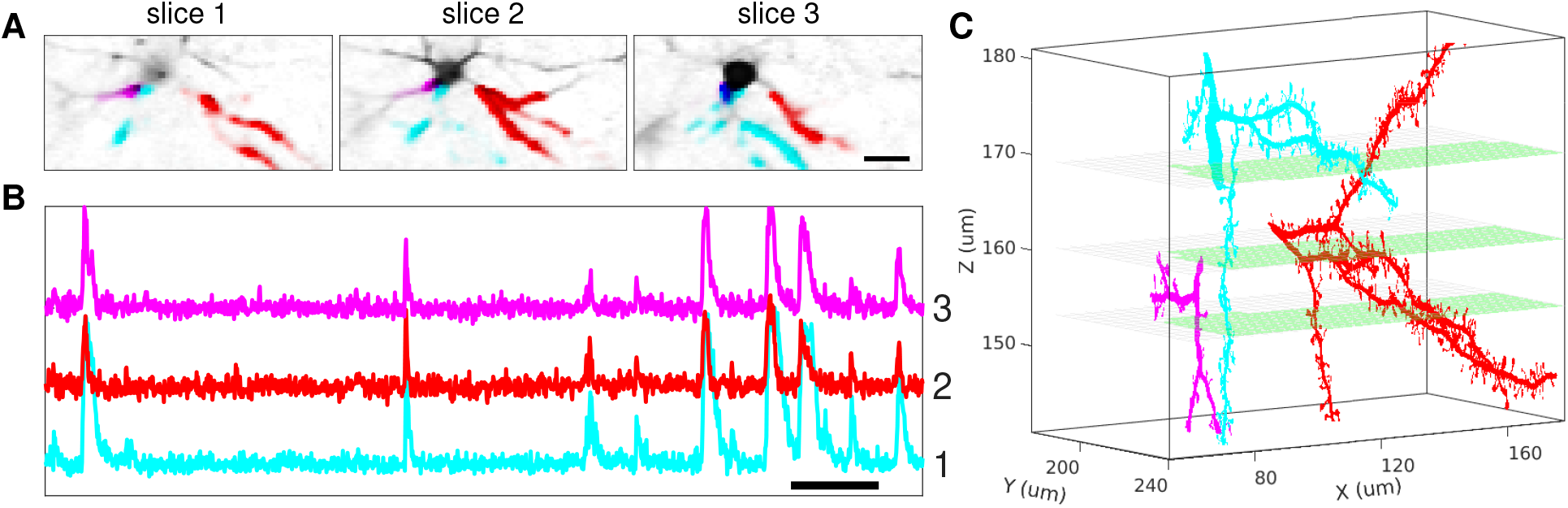
EASE can enable joining of split EM segments. (**A**) spatial footprints of three temporally correlated neurons (magenta, cyan, and red) and the correlation image (gray) between the raw video *Y* and the mean of their estimated traces *c_i_*. Green rectangle indicates the boundaries of the EM volume; scale bar: 20 um. (**B**) estimated temporal components of the three selected neurons (scale bar: 10 seconds); note the strong correlation. (**C**) The EM meshes of the three neurons. (The green areas indicate the scanning planes.)

### 2.3 Joint processing of multiple functional scans

So far we have focused on extracting neural components from a single functional scan. One of the major advantages of joint CI-EM data is that we can fuse information from multiple functional scans, by taking advantage of the fact that a given three-dimensional EM segment may extend over multiple scans, allowing us to match the functional signals extracted from each scan back to the same neuron. Figure 7 illustrates this idea: panel A displays a single three-dimensional EM segment, with the corresponding footprints *p_i*_* computed from this segment shown in panel B. Panel C displays the functional spatial components *a_i_* extracted from four separate functional scans, matched to the EM footprints *p_i*_* shown in panel B. Panel D displays the corresponding temporal components *c_i_* extracted from the four scans, and panel E shows that the visual direction tuning curve extracted from each temporal component is consistent across these scans, providing a useful secondary check on the quality of the matchings computed here.

**Figure 7:**
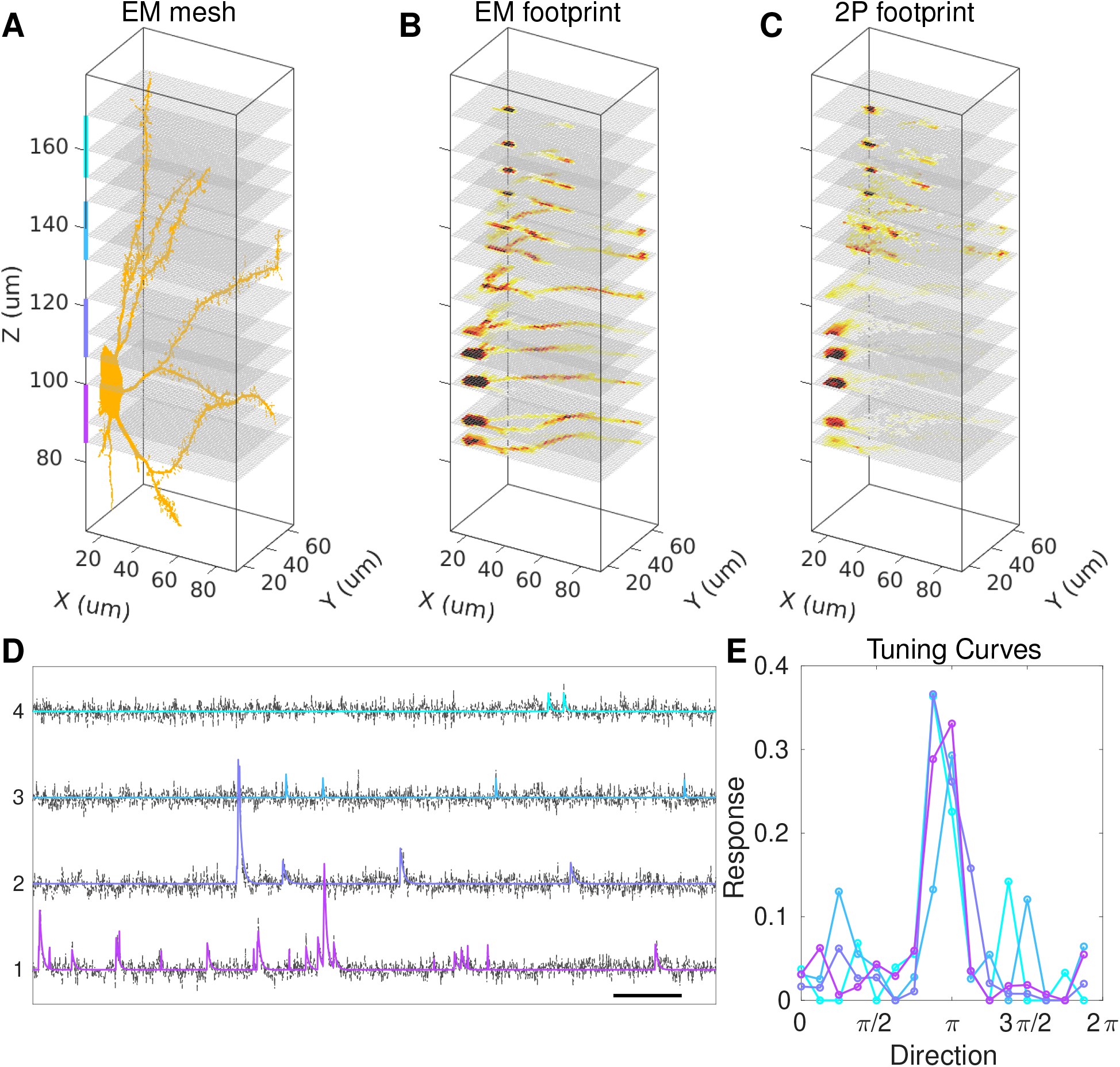
Merging the activity from a single neuron over four separate functional imaging scans. (**A**) High-resolution EM meshes of an example neuron in the whole 3D volume. Vertical colored bars group the three planes imaged within the same scan. (**B**) EM footprints *p_i*_* on the imaging planes of the four scans. (**C**) The extracted spatial footprints of the neuron on the same imaging planes. (**D**) The noise-normalized temporal traces *c_i_* of this neuron imaged over the four scans (scale bar: 10 seconds). (**E**) The direction tuning curves extracted from this neuron in each of the four scans.

### 2.4 Comparison against constrained nonnegative factorization without EM constraints

How well can CNMF methods without additional EM structural constraints recover the components extracted here? To address this question, we ran the pipeline from (Buchanan et al., 2018) on a single functional scan, and compared the resulting spatial and temporal components to the components output by the full multi-scan EASE pipeline described above. (Other demixing pipelines led to qualitatively similar results; data not shown.) Figure 8 compiles the results: we see that the pipeline from (Buchanan et al., 2018) recovers the brightest, highest-PNR components well, but misses many dimmer components; EASE recovers about twice as many components (210 vs 107). Importantly, we observe minimal dependence between the PNR and the degree of visual tuning of the recovered components, as measured by the visual direction sensitivity score described in Section 4.11.

**Figure 8:**
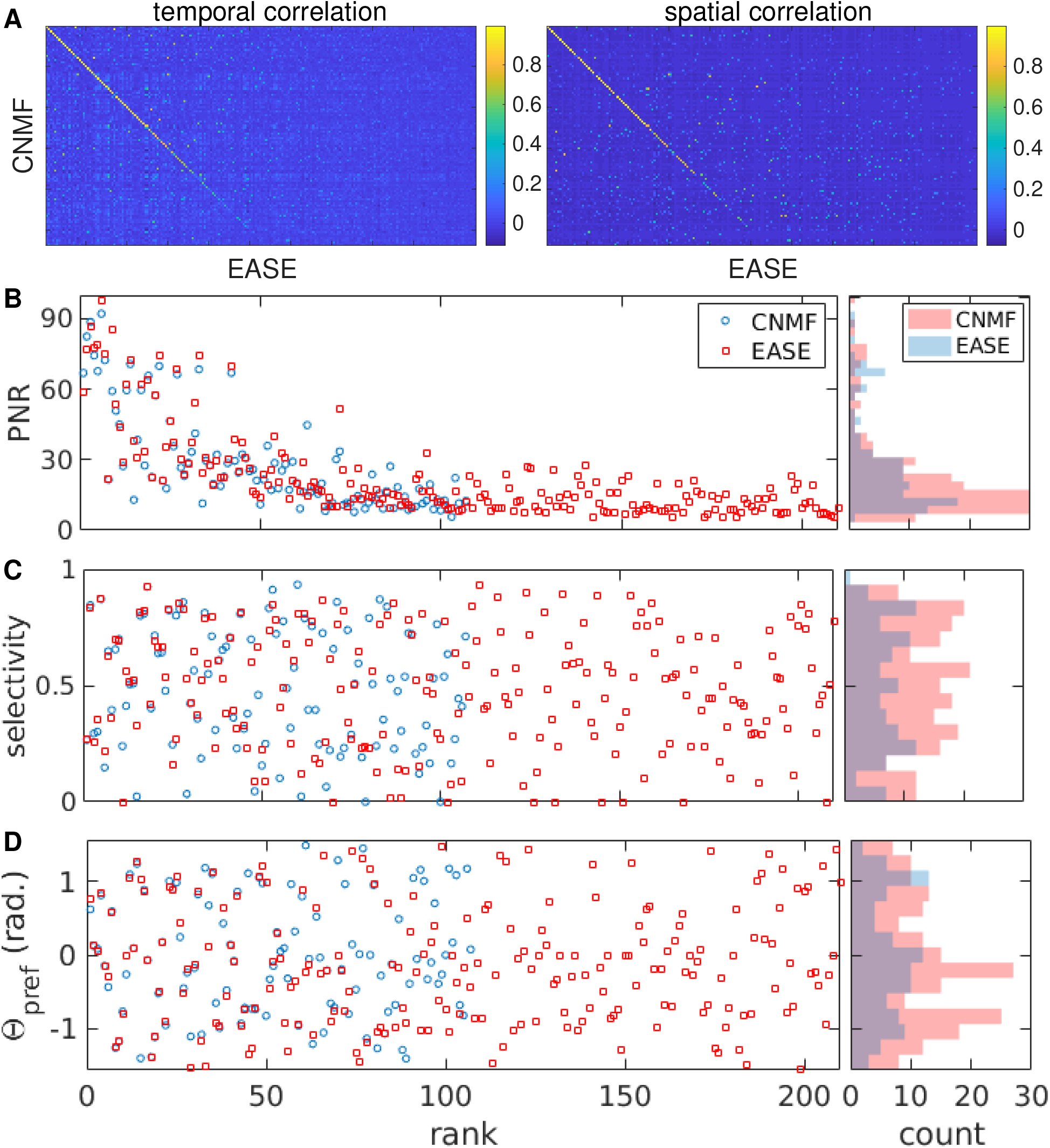
Comparison between the results of EASE and a demixing pipeline without access to EM structural prior information. (**A**) spatial and temporal pairwise correlations between neurons extracted using EASE and the CNMF-based pipeline from (Buchanan et al., 2018), respectively. Components have been ordered to greedily maximize the diagonal of the left panel, to perform a simple matching. EASE extracts 210 components from a single functional scan here, compared to 107 without EM structural prior information. (**B**-**D**) PNRs (**B**), visual direction tuning scores (**C**; Section 4.11) and preferred directions (**D**) for all components, using the same ordering as in (**A**). The right panels show the corresponding histograms. Note that visual direction sensitivity does not depend strongly on PNR.

One possibility is that these new low-PNR components recovered by EASE might be redundant with the higher-PNR components already recovered by standard pipelines. The results in Figure 8A+B argue against this possibility, since the correlations in the low-PNR components do not seem to be systematically stronger than those in the high-PNR components. To examine this possibility further, we performed an additional decoding analysis, attempting to map the observed neural activity into an estimate of the visual stimulus orientation *θ* on each trial. We find that decoding using the EASE output leads to consistent, statistically significant improvements in decoding error compared to the localNMF outputs, with mean absolute error decreasing from 21 ± 0.8 degrees (using the localNMF output) to 16 ± 0.7 degrees (using the EASE output). Thus, in summary, EASE extracts a large number of low-PNR but visually-tuned components that are currently difficult to extract without the side information provided by the EM constraints in EASE; moreover, these low-PNR components carry a significant amount of visual information that is not completely redundant with the high-PNR components extracted by standard CNMF-based pipelines.

### 2.5 Using the EASE output to simulate spatial components extracted from 2p imaging and train a spatial component denoiser

We expect that one critical long-term application of the results presented here will be in scoring calcium imaging analysis pipelines: i.e., if we input the raw 2p imaging data (without knowledge of the EM components) into an analysis pipeline and the output closely matches the EASE output, then the pipeline would achieve a good score. The availability of this EM-constrained scoring mechanism will help guide further improvements to analysis pipelines in the near future.

In the shorter term, however, we can already make use of the EASE output to improve analysis pipelines. As a first illustration, in this section we demonstrate how to use the EASE output to generate realistic simulated data and to train a neural network (NN) to denoise the spatial components *a_i_*. This denoiser in turn can be dropped into any analysis pipeline to improve signal extraction.

The starting point of any NN estimation procedure is to gather a large set of training data. Gathering labeled training data is often a slow and labor-intensive process. However, we have access here to a large dataset of high-resolution three-dimensional EM component shapes; if we can take multiple slices to convert these three-dimensional EM shapes into corresponding two-dimensional shapes at 2p resolution, then we would have a strong dataset for training NNs, without requiring any additional hand-labeling. Following this logic, we construct our NN training set in two steps. First, we fit a model that maps three-dimensional EM shapes into corresponding estimates of the two-dimensional spatial components *a_i_*. Then we applied this model to slice each EM component in multiple planes into simulated *a_i_* images, and used the resulting large set of simulated *a_i_* images as our training set.

In the first step, we experimented with multiple versions of this EM-to-*a_i_* model. Our baseline model simply outputs *p_i_* as the estimate for *a_i_*; recall that this model assumes the brightness is uniform across all EM voxels, and just applies a convolutional point-spread function (psf) to map the EM component into the imaging plane to generate *p_i_*, as illustrated in Figure 7B. In practice, the resulting *p_i_* tends to have brighter dendritic processes than the corresponding estimated *a_i_*; this effect is clearly visible in Figures 3 and 7.

To improve our predictions beyond this baseline *p_i_* model, we included a term to modulate the brightness of each voxel before applying the 3d-to-2d psf transformation. We experimented with three such models: the brightness could be a function of the distance from the center of the soma, or it could be a function of the distance from the surface of the cell, or the brightness could depend on both of these factors. We found that incorporating a spatial modulation term led to significantly improved prediction accuracy; all three versions of the model provided similar improvements. Specifically, the model learned that voxels near the surface of the cell tend to be dimmer than voxels further into the interior of the cell; moreover, voxels near the nucleus tend to be relatively dim (i.e., the models learn to predict the classic “doughnut” shape of cytosolically-labeled somas). See Figure 9; full model details are provided in the Methods section.

**Figure 9:**
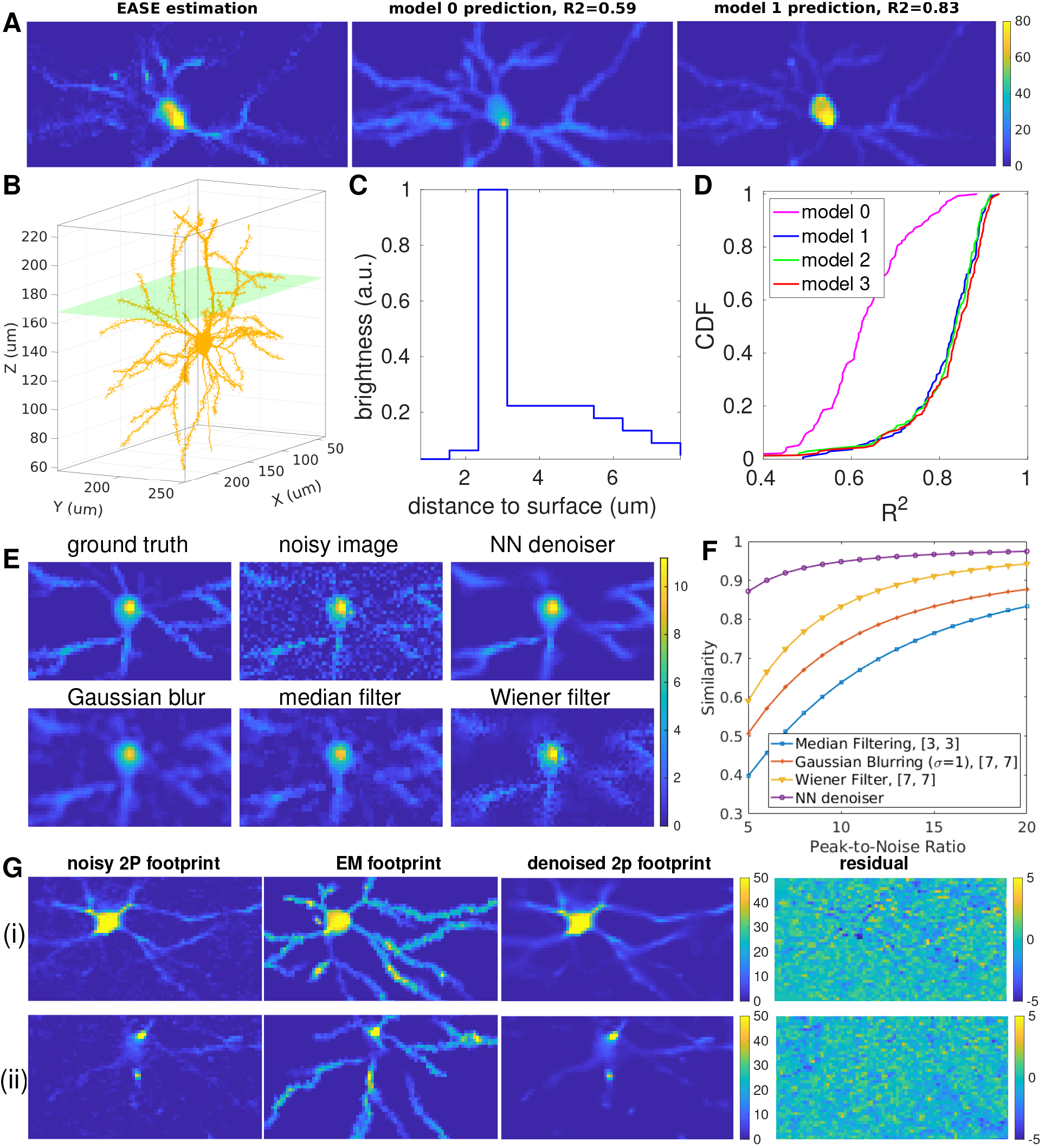
The EASE output can be used to fit a model to predict a neuron’s two-dimensional 2P *a_i_* image from the corresponding three-dimensional EM segment, and these predictions in turn provide clean training data for a neural network (NN) denoiser that can enhance 2P analysis pipeline estimates. (A) The *a_i_* estimated by EASE from CI data next to the *a_i_*’s predicted from its EM segment (B). Model 1 assumes that voxel brightness is a function of the distance to the cell membrane surface; model 0 assumes a uniform brightness per voxel, and leads to a worse prediction (with overly-bright dendrites). The green plane in (B) indicates the 2P imaging plane. (C) The estimated relationship between the brightness of each voxel and its distance to the cell surface; note that surface voxels are relatively dim, as are voxels that are furthest from the surface (i.e., nearest to the nucleus). (D) Cumulative distribution function of the *R^2^* for different model predictions on 135 test images. (Models 2 and 3 incorporate dependence of the voxel brightness on the distance to the soma; see appendix for details.) Models 1-3 achieve similar accuracy; all of these models significantly outperform the simpler model 0. (E) Example simulated ground truth image *a_i_*, along with the simulated noisy observation and output of different baseline image denoisers. The trained NN denoiser achieves the qualitatively best denoising results, as quantified further in (F), which tabulates denoiser performance (measured as cosine similarity between ground truth vs recovered test image; *N* = 24 images) under multiple signal-to-noise levels. (G) Performance of the trained image denoiser on two example test spatial components *a_i_*. The corresponding EM footprint is also shown for illustration purposes (but is not provided to the NN denoiser); note that the residual (the noisy observed *a_i_* minus the NN denoiser output) is relatively featureless, indicating that the denoiser error is dominated by noise, not model bias. (Note that all colorbar units in this figure are arbitrary, since *a_i_* is only defined up to a scale factor.)

One useful application of the resulting large pool of simulated *a_i_* images is to generate simulated 2p calcium imaging videos, to test the accuracy of the EASE pipeline (and potentially other pipelines). (See (Charles et al., 2019) for a detailed biophysical simulation, in which the neural shapes are generated by a random process instead of deriving the shapes from three-dimensional EM segments, as we do here.) See Appendix section 4.14 for details; we find that the EASE pipeline achieves high accuracy on realistic simulated data.

Finally, we trained a NN denoiser using simulated *a_i_* images drawn from the EM-to-*a_i_* model. Specifically, the NN was trained to take noise-corrupted *a_i_* images as input and to output the best estimate of the original clean *a_i_* image. Figure 9E-G illustrate the denoiser at work on several test images, along with comparisons against several simpler image-processing baselines. The NN learns to suppress speckle noise across a wide range of signal-to-noise regimes, while avoiding oversmoothing small dendritic processes much more effectively than do the generic image-processing baselines.

## 3 Discussion

We have introduced EASE, a method to fuse calcium imaging (CI) and electron microscopy (EM) data. In EASE, dense reconstruction of EM anatomy is exploited to enable dense, constrained extraction of functional CI signals. The resulting spatial components display a large degree of overlap (c.f. Figs. 4 and 5A), emphasizing the importance of *demixing* these functional signals (i.e., modeling the signal within each pixel as arising from potentially multiple neurons), rather than simply *segmenting* these images (i.e., constraining each pixel to be “owned” by at most a single neuron). Furthermore, in the dataset examined here, about 6× as many dendritic components as somatic components were extracted by EASE. Thus, restricting attention to somatic components may leave significant information behind in CI recordings. In addition, we find that EASE recovers a significant number of dim but visually-tuned components that are not recovered by standard constrained non-negative matrix factorization (CNMF) methods; thus there may be room to further improve existing CI analysis pipelines, using these combined CI-EM data as a valuable ground truth benchmark.

EASE outputs estimates of the temporal activities and spatial shapes of the neurons visible in the field of view, matched to their corresponding EM segments. We are releasing these components publicly at this **site**. As emphasized in (Paninski and Cunningham, 2018; Soltanian-Zadeh et al., 2019), we have to date lacked ground truth datasets for the CI demixing problem. Since good training sets are a critical ingredient in the “secret sauce” for accelerating progress in data science (Donoho, 2017), we hope that this new EM-constrained gold standard dataset will help spur further development of improved algorithms for CI analysis that can be applied in the vast majority of cases where no EM side information is available to further constrain the estimates. The neural network denoiser illustrated in Figure 9 is a first step in this direction.

Finally, it is worth noting that a version of the approach developed here should be applicable to other dense micro-anatomical approaches that are currently in development (Alon et al., 2018; Gao et al., 2019): given a three-dimensional segmentation of the anatomical channel, we can import the resulting segments, compute the footprints *p_i*_*, and use these to constrain the spatial footprints *a_i_* We hope that these tools will enable the interrogation of dense structure-function relationships in a wide variety of neural circuits in the coming years.

## Acknowledgments

We thank Sebastian Seung’s group for generously sharing their EM segmentation prior to publication. We thank E.K. Buchanan for useful conversations, and J. Zhao for helping with the design of Figure 1.

## Author Contributions

PZ and LP developed the EASE method and wrote the manuscript, with input from the other authors. PZ wrote the EASE software and performed all analyses described here, with supervision from LP. DZ and IK aided in the pipeline comparisons described in Section 2.4. DZ aided in the decoding analysis described in Section 2.4. AP aided in the analyses summarized in Figure 9. JR, EF, DVY, PGF, and AST collected the CI data. AB, JB, DB, GM, RT, NC and CR oversaw the collection of the EM data. SD, DI, KL, RL, TM and JW reconstructed the EM data and registered it to the 2p stack data. JR coordinated the collaboration.

## Declaration of Interests

The authors declare no competing interests.

## 4 Methods and Materials

### 4.1 Key Resources Table

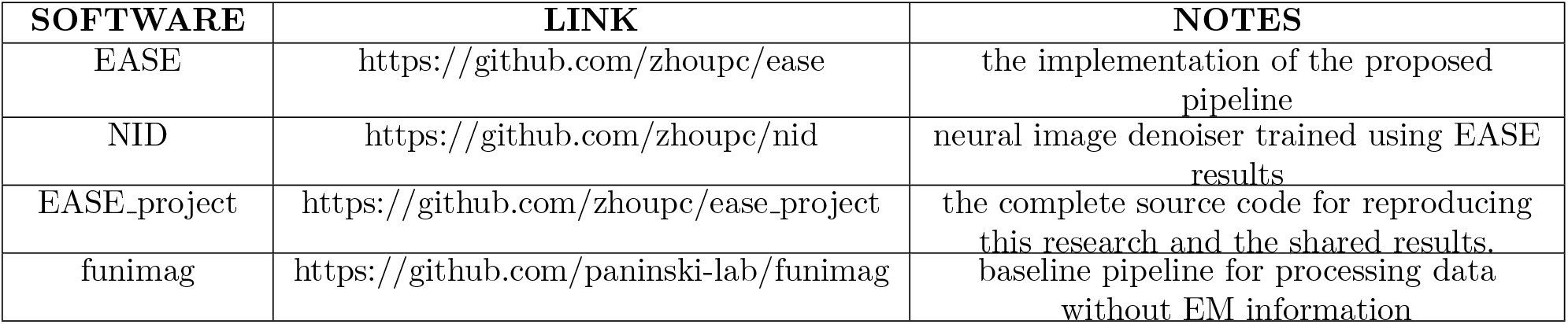

### 4.2 Overview

We begin with the CI data *Y*, the three-dimensional EM segmentation, and a two-photon structural scan of the same volume (which helps to spatially align the CI and EM data). We co-register the EM and CI data and then compute the intersection of the three-dimensional EM data with the two-dimensional CI scan, then spatially downsample and blur the EM data to match the two-photon resolution and obtain a set of two-dimensional EM “footprints” {*p_i_*} (see details below). Next, we greedily initialize active neurons using these footprints to index into the CI data *Y*. We compute iterative CNMF updates using these initialized components, enforcing the constraint that each functional spatial component *a_i_* must lie within the support of the corresponding EM footprint *p_i*_* (*i*^*^ is the index of the EM segment that matches *a_i_*). Then we iterate, adding more components and running CNMF updates until no further good components can be added. We have also developed several diagnostics for component quality that can be used to discard bad components within this loop.

Below (section 4.3 - 4.10) we provide full details on each of the above steps.

### 4.3 Data description

The analyses described here are based on three types of data acquired in sequence from the same volume:

- Structural volume image: imaging was performed in a triple-transgenic CamK2a-Cre/Camk2a-tTA/Ai93 mouse (Madisen et al., 2015). In this mouse, GCaMP6f expression was restricted to pyramidal cells, and the Tet enhancer system produced strong expression of the fluorophore, resulting in higher SNRs at lower imaging powers. We obtained a 3D image of a 400 × 400 × 310 um^3^ volume using conventional two-photon (2P) laser scanning microscopy. After pixelization this data set has dimensions 512 × 512 × 310.
- CI data *Y*: we took two-dimensional functional scans of the same FOV as the (3D) 2P structural volume described above. The spatial resolution of each functional scan was half that of the 2P structural volume per plane (256 × 256). The measured point-spread function was approximately 0.5 × 0.5 × 3 um in x, y, and z; pixel pitch in x and y were approximately 1.6um. (Importantly, this resolution was sufficiently fine to enable us to record functional activity from many distinct apical dendrites traversing the imaging volume.) We took 9 scans in total; each scan contains three z planes and all three planes were imaged nearly-simultaneously, with full acquisition over all three planes at 14.8313 frames per second.
- High-resolution EM segments: an electron microscopy data volume was acquired within the 2P structural volume, then segmented computationally by Sebastian Seung’s group at Princeton University (see (Dorkenwald et al., 2019) for full details). In this work, we used 109084 segments from the EM reconstruction of a volume of 196 × 129 × 40 um^3^ (note that this is a significantly smaller volume than was imaged in the 2P structural scan). These segments were saved in the format of triangular meshes and the mesh coordinates were registered to the 2P structural volume image.

### 4.4 Preprocessing

**Co-registration of EM data and 2P stack data**: We found 189 cell bodies in both the 2P stack and EM data, computed their centroids, and then fit a regression model to obtain an affine transformation model between the 2P and EM data (full details will be described in a manuscript in preparation).

**Voxelization of EM segments**: The segmented EM components are saved into meshes. For the analysis below, we need voxelized representations to obtain the spatial support of each segment within each imaging plane. We used *polygon2Voxel*^1^ for voxelizing the surfaces, then fill to obtain the entire 3D segments. The voxelized EM components can be represented as a binary multidimensional array *M* ∈ {0,1}^512^×^512^×^310^×^*K_em_*^. Note that *M* is a huge but very sparse array; thus we only need to save the indices of nonzero voxels.

**Cropping the FOV**: The CI FOV is larger than the EM volume. To crop the CI FOV to match the EM volume, we projected the EM volume to the x-y plane and chose a slightly larger rectangle (5 more pixels surrounding the borders) around this.

**Motion correction of CI data**: Raster artifacts from resonant scanning were corrected post-hoc, and X/Y subpixel motion correction was performed as in (Reimer et al., 2014).

**Co-registration of CI data *Y* and 2P stack data**: We register the averaged motion-corrected functional scan data onto the 2P anatomy stack via cross-correlation.

**Projection of the EM segments intersecting with one scanning plane**: Our model is based on constraining each functional neural shape to match the shape of the corresponding EM segment. The EM segments are three-dimensional but the functional imaging scans are two-dimensional, so we need to compute the intersection of the EM components with the functional imaging planes. In addition, we need to account for the fact that the spatial resolution of the two-photon functional imaging data is much lower (particularly in the axial direction) than the EM resolution. In short, we would like to model what each individual EM component would look like if imaged by two-photon scanning in a given plane (assuming that the calcium indicator has a uniform brightness in each cell). To obtain these shapes, we selected all pixels in the EM mask matrix M that were near the imaging plane, then applied a Gaussian blurring in z and spatial downsampling in xy to emulate the point-spread function of the 2P scanning microscope, which is extended in z. We used a Gaussian width of *σ* = 8 *μm* and included EM pixels within a symmetric 32 *μm* window around the imaging plane. The resulting matrix of projected EM components is denoted *P* = [*p*_1_,*p*_2_,⋯,*p_K_em__*]. See Figure 2 for some examples.

**Noise-normalization**: We normalize each pixel in *Y* so that the noise level (estimated via the power-spectrum density method described in (Pnevmatikakis et al., 2016)) is constant across pixels.

### 4.5 Model

See Table 1 for a summary of the variables used below.

**Table 1:** List of variables. For the dataset analyzed here, *d*_2*p*_ = 512 × 512 × 310 is the number of voxels for the full 2P stack data. *K_em_* = 109084 is the number of EM components. d =58 × 129 × 3 is the number of pixels for one scan of the cropped 2P video data; *T* = 27100 is the number of frames; K is the number of extracted neural components, which is around 150; *N* = 3 is the rank of the background B^f^.

| Name | Description | Domain |
| --- | --- | --- |
| $M$ | spatial mask of all EM segments | $\{0, 1\}^{d_{2p} \times K_{em}}$ |
| $Y$ | motion corrected video data | $\mathbb{R}^{d \times T}$ |
| $A$ | spatial components of neurons | $\mathbb{R}_+^{d \times K}$ |
| $P$ | spatial footprints of EM segments | $\mathbb{R}_+^{d \times K_{em}}$ |
| $C$ | temporal components of neurons | $\mathbb{R}^{K \times T}$ |
| $B$ | background | $\mathbb{R}^{d \times T}$ |
| $E$ | noise | $\mathbb{R}^{d \times T}$ |
| $D$ | spatial components of the background | $\mathbb{R}^{d \times N}$ |
| $F$ | temporal components of the background | $\mathbb{R}^{N \times T}$ |
| $\mathbf{b}_0$ | temporally-constant baseline per pixel | $\mathbb{R}^d$ |
| $B^f$ | temporally-fluctuating part of the background | $\mathbb{R}^{d \times T}$ |

We used the constrained nonnegative matrix factorization (CNMF) framework (Pnevmatikakis et al., 2016; Zhou et al., 2018) for modeling the calcium imaging data:

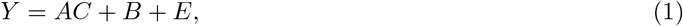

where *Y* represents the video data as a matrix (Pnevmatikakis et al., 2016), *A* = [*a*_1_,*a*_2_,…,*a_K_*] contains the inferred neuron shapes, and *C* = [*c*_1_,*c*_2_,…,*c_K_*]^*T*^ contains the inferred calcium traces of all neurons. Following (Zhou et al., 2018), we decompose the background term *B* into a constant baseline term and a fluctuating term, *B* = *b*_0_1^*T*^ + *B^f^*. We find that the fluctuating term *B^f^* in the small FOV examined here is well-modeled as a low-rank matrix:

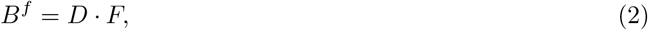

where *D* and *F* have rank *N* = 3 here. Following (Vogelstein et al., 2010; Pnevmatikakis et al., 2016; Friedrich et al., 2017b; Zhou et al., 2018), we model the calcium dynamics of each neuron *c_i_* with a stable autoregressive (AR) process of order *p* = 1,

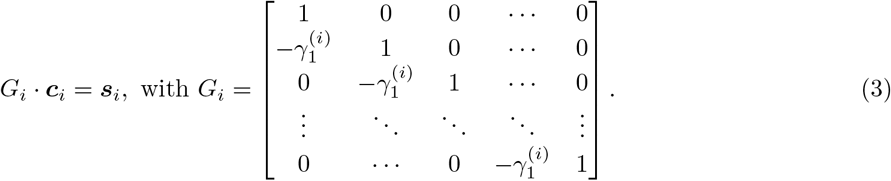

where *s_i_*(*t*) ≥ 0 is the number of spikes that neuron fired at the *t*-th frame. The AR coefficients 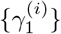 are different for each neuron and they are estimated from the data.

The novel part of the model here is that in (1), each *a_i_* has its EM counterpart *p_i*_*. Recall that *p_i*_* models the two-photon appearance of cell i assuming uniform calcium indicator brightness throughout the cell; thus the support set of *a_i_* (i.e., the nonzero pixels in *a_i_*) should be contained within the support of *p_i*_*, but due to non-uniformities in indicator brightness *a_i_* and *p_i*_* will not match exactly.

### 4.6 Model fitting

After initialization of the components *c_i_, a_i_*, along with the corresponding EM support sets *p_i*_* (see the next section for initialization details), we minimize the residual sum of squares (RSS) given multiple constraints for estimating all model variables as a single optimization meta-problem

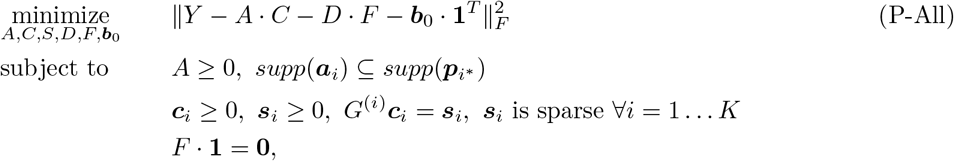

where 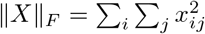 denotes the Frobenius norm of the matrix *X* and *K* is the number of extracted neural components. Similar to (Zhou et al., 2018), we divided the nonconvex problem (P-All) into three simpler sub-problems:

**Algorithm 1.**
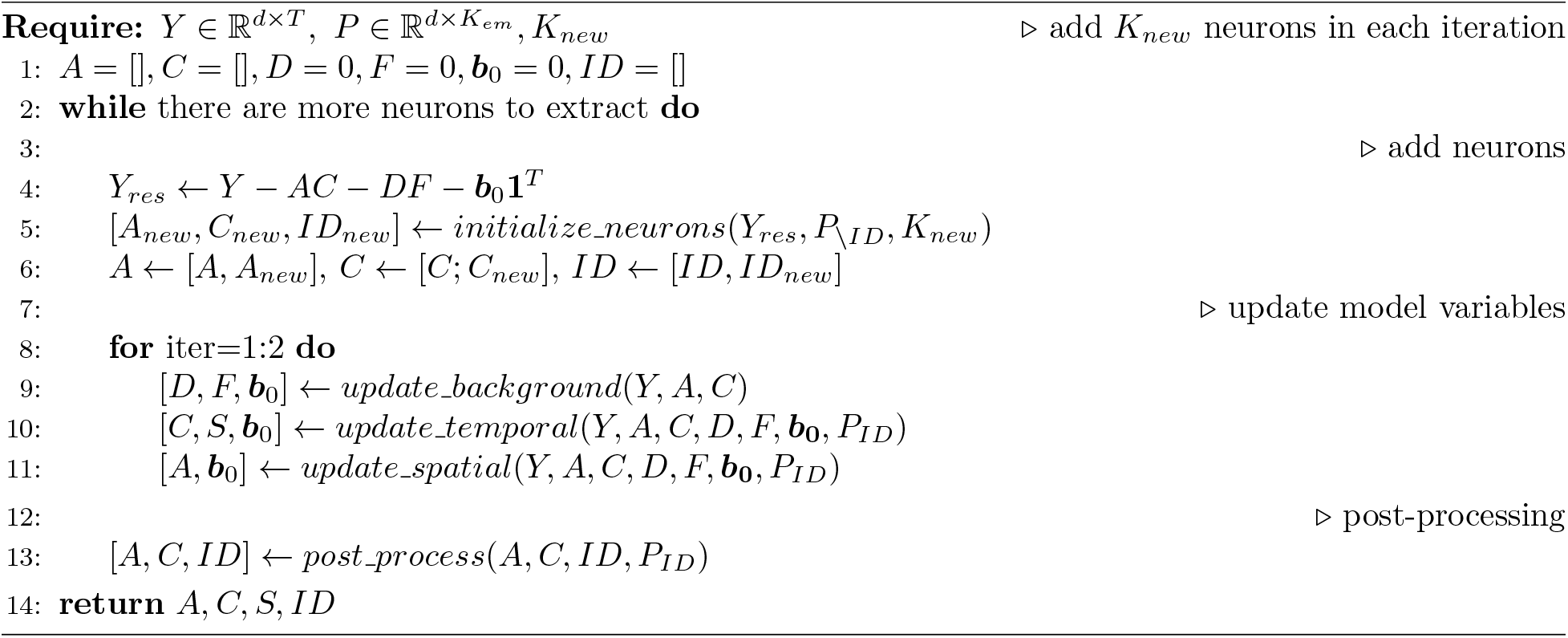
EASE Pipeline

Estimating *A, b*_0_ given 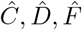

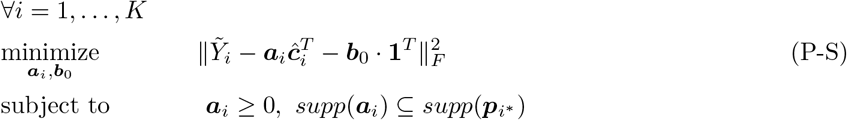

Estimating *C, b*_0_ given 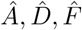

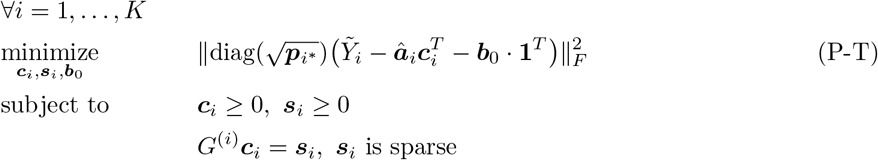

Estimating *D, F, b*_0_ given 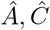

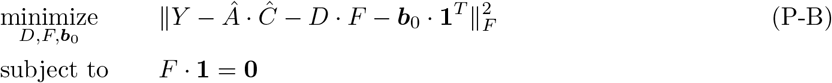

In both problems (P-S) and (P-T), 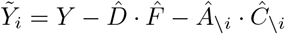 denotes the residual after subtracting the fluctuating background and the spatiotemporal signal of all neurons except the *i*-th neuron. The objective function in the problem (P-T) includes an additional upweighting term 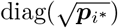; this upweights the squared residuals at each pixel by a factor proportional to *p_i*_*. We found that this upweighting led to improved signal recovery, particularly in early iterations when a number of visible neural components remain in the residual; we will discuss this issue in more depth in the next section.

We perform updates to solve each of these three sub-problems in alternating fashion. Details are summarized in Appendix (5.4) and Algorithm (3). After convergence (typically in just a couple iterations) we perform a manual or automatic post-processing step (to potentially remove any false positives and correct any clearly mistaken matches) and then initialize more components, and iterate. See Algorithm 1 for an overview, and further details in the following sections.

### 4.7 Initializing neural components constrained by EM segments

As emphasized in previous work (Pnevmatikakis et al., 2016; Zhou et al., 2018), due to the nonconvexity of the constrained NMF problem, good initializations are critical — especially so here, since poor initializations of the spatial components *a_i_* may lead to poor matches with the EM components *p_i*_*, leading to poor local optima.

Our approach is to start by finding EM components that are likely to correspond to “good neurons” in *Y*, and use these to initialize the components *a_i_* and *c_i_*. Then we subtract the estimated spatiotemporal activity of these initialized neurons from *Y* and iterate. This greedy algorithm provides a scalable approach that simultaneously extracts neurons from *Y* and finds matches with the corresponding EM components.

How do we find a good set of EM components that contribute significantly to *Y*? The simplest way to proceed would be to simply project the data *Y* onto the EM components *p_i*_*, and initialize components with the biggest projections. Unfortunately we find that this does not work in practice, because a *p_i_* that happens to spatially overlap with a bright component in *Y* can “steal power” from this component, leading to a mistaken initialization that may be difficult to correct downstream. (Recall that *p_i_* is a blurred, downsampled projection from the original high-resolution three-dimensional EM volume — where no components overlap spatially, by construction — into the lower-resolution functional imaging plane, where multiple components overlap spatially.)

Therefore we need an improved approach for initializing functional components *a_i_* from the EM components *p_i_*. The approach described below is a critical technical contribution of this paper. See Algorithm 2 for a summary.

**Algorithm 2.**
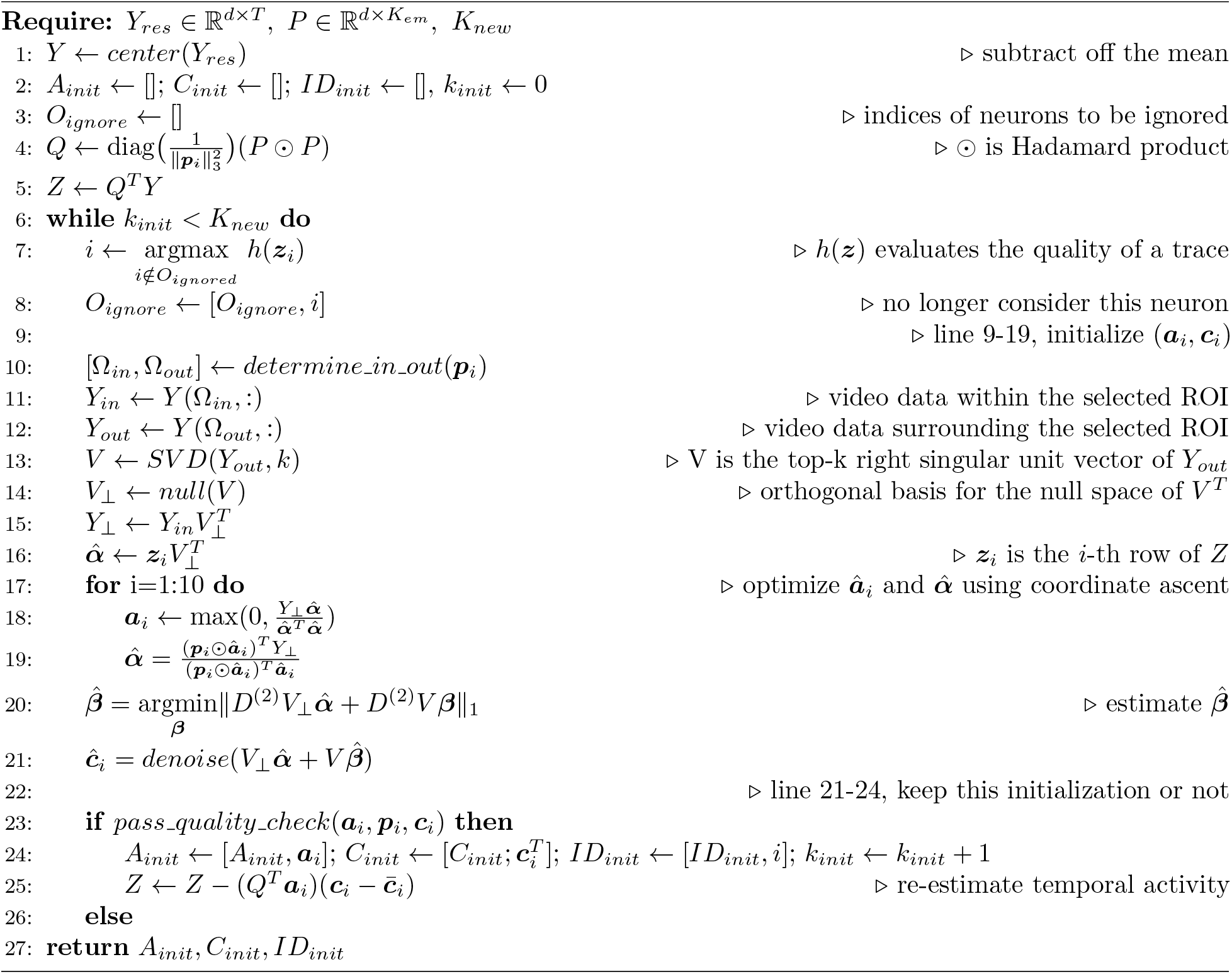
Initialize neurons using EM masks

#### 4.7.1 Sorting the EM components

Since we have roughly 100 × more EM components than functional components in *Y* (compare Figure 2 to S1 Video), we need a quick rough method to order the EM components *p_i_* which are likely to be most active in the functional calcium video data. Given a method for rapidly converting *p_i*_* into a corresponding crude initial estimate of (*a_i_, c_i_*) (to be defined below), we found the following rough score to be useful for this task:

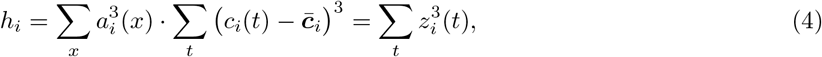

where 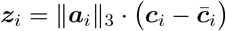 and 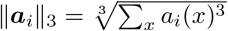 is the 3-norm of the vector *a_i_*. (We used the 3-norm here because we found that it was more effective than e.g. the 2-norm in capturing non-negative, positively-skewed signal in the components *a_i_* and *c_i_*.) Components with larger values of h¿ are more likely to contain useful activity.

To rapidly extract a coarse initial estimate from *p_i*_*, we set *a_i_* = *p_i*_* and then estimate *c_i_* by minimizing the objective function in problem (P-T) while ignoring all constraints:

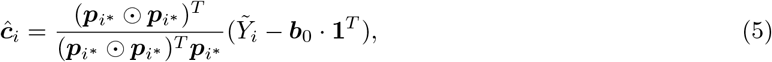

where 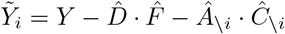 Then we can further simplify the form of *z_i_* as

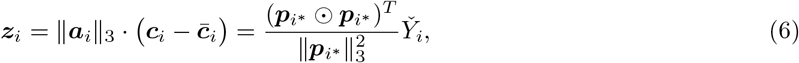

where 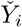 is centered to have a temporal mean of 0.

Thus we can utilize simple matrix computations to quickly estimate all *z_i_* at once and then calculate the corresponding *h_i_*. Every time we initialize a neuron 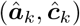, we can easily update the remaining *z_i_* values:

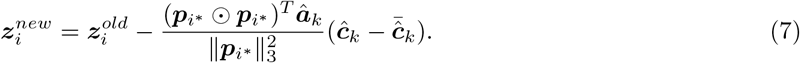

#### 4.7.2 Initializing a single neuron given the EM footprint

The raw data *Y* contains the summed spatiotemporal activity of many neuronal components; thus initializing the spatial and temporal components of one neuron using a given EM footprint *p_i*_* requires the successful removal of contaminations from other neurons and background signals that spatially overlap *p_i*_*.

The simplest approach to initializing *a_i_* and *c_i_* from *p_i*_* would be to initialize *a_i_* = *p_i*_* and then compute a semi-NMF factorization: 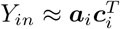, where *Y_in_* denotes the mean-subtracted video data *Y* restricted to the support of *p_i*_*, and we restrict *a_i_* ≥ 0. However, we found that this approach was not sufficiently robust to contamination in *Y_in_* from other spatially overlapping cells; Figure S1 below provides an example.

To develop a more robust approach we utilize the spatial support of *p_i*_* more explicitly. Denote the pixels within *p_i*_* as Ω¿_n_ = supp(*p_i*_*), and the surrounding set of pixels (within a distance of 8 pixels) as Ω_out_. We observe that large contaminations often span across both Ω*_in_* and Ω*_out_*, while the spatial range of the targeted neuron is by construction constrained within Ω*_in_*. Thus we model the (mean-subtracted) fluorescence signal in Ω*_in_* as

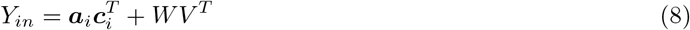

where *a_i_* ≥ **0**, *V* is the matrix of the top-k right singular unit vectors of the fluorescence data of pixels in Ω*_out_* (in practice we find *k* = 10 works well to summarize the activity in Ω_*out*_), and *W* is a matrix of weights applied to *V*, with a different weight vector for each pixel in Ω*_in_*. Thus the role of *WV* is to help explain away contaminations in Ω*_in_* from background fluctuations and overlapping neurons (see Section 5.5 for a detailed derivation of Eq. 8).

Fitting this model requires estimating *a_i_, c_i_*, and *W*. We first compute an orthogonal basis for the null space of *V^T^* as *V*_⊥_, and decompose *c_i_* as *c_i_* = *V*_⊥*α*_ + *V^β^*. Then Eq. (8) becomes

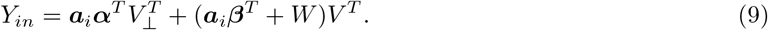

One natural approach for fitting *a_i_, α, β*, and *W* is to minimize the weighted residual sum of squares (wRSS),

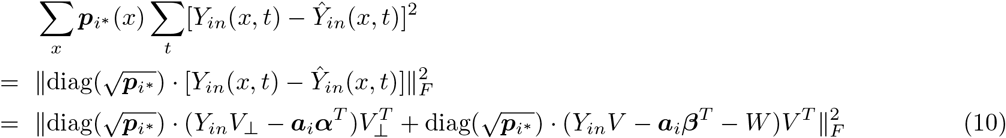

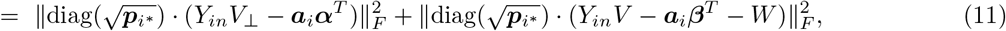

where 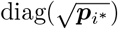 weights the residual of different pixels according to the selected neuron’s EM footprint^2^; in practice, we found that this weighting can further reduce the neighboring neurons’ contamination, as discussed below. Since W is unconstrained, the second term in the right hand side of Eq. (11) can always achieve its minimum of 0 by letting 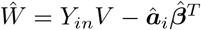. Thus minimizing the wRSS is equivalent to minimizing 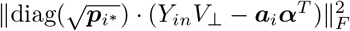. However, solving this new optimization problem

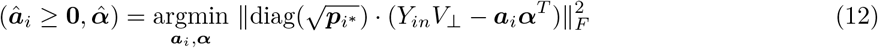

only yields estimates of *a_i_* and *α*, providing no information about *β*.

We resolve this issue by exploiting the temporal smoothness of *c_i_*, i.e., we estimate *β* by minimizing 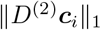 to enforce its smoothness, where *D*^(2)^ is the second order discrete difference operator of order 2. So given 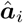 and 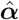, the estimated 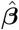 is

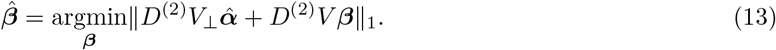

In summary, we first estimate 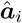 and 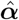 by solving problem (12), and then estimate 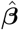 by solving problem (13), to obtain 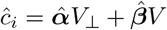. We use block coordinate descent (Cichocki et al., 2007) to optimize *a_i_* and *α* in (12); we find 10 iterations suffice. We optimize *β* in (13) with subgradient descent; we use backtracking line search to determine step sizes (Boyd and Vandenberghe, 2004). Empirically we found that our method significantly outperforms the naive semi-NMF method in initializing *a_i_* and *c_i_*; see Figure S1 for an illustration.

Finally, before adding the *â_i_* and *ĉ_i_* computed above to our set of components *A* and *C*, we run a simple quality check on the component. Occasionally the selected *p_i*_* does not contain a prominent fluorescence signal. This typically leads to an estimated *â_i_* that matches *p_i*_* poorly. To detect these cases we compute the cosine similarity between the initialized *â_i_* and *p_i*_*, discarding initializations with similarities smaller than 0.55.

### 4.8 Evaluating the quality of cell matching

EASE assigns an EM match to each neuron extracted from CI data during the initialization step, but not all of these preliminary assignments are correct. For each extracted component, we would like to know how well this neuron matches to each EM component and how confident we should be about the current match. Hence we define two metrics:

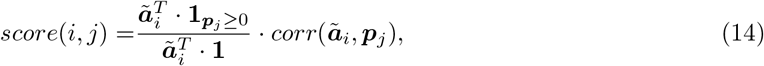

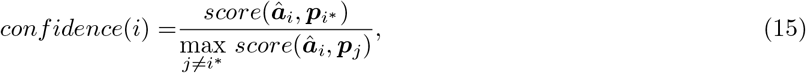

where *ã_i_* is the unconstrained estimation of *a_i_* in problem (P-S); recall that *i*\* denotes the current match to the *i*-th neuron. The score above combines a correlation between *ã_i_* and *p_j_* with a term that measures the fraction of *â_i_* contained in the support of *p_j_*: we found that combining these two terms led to a score that better matched the visual overlap between *â_i_* and *p_j_* (compared to using either the correlation or support overlap alone). The “confidence” in the second line above simply computes the score ratio of the assigned match and the best alternative match.

### 4.9 Jointly extracting neurons from multiple functional imaging scans

In this dataset multiple functional scans were acquired at different *z* depths at different times. When run on a single functional scan, EASE outputs the inferred two-dimensional spatial components *a_i_* from this scan, along with their matched three-dimensional high-resolution EM segments. By tracing the EM reconstruction from one scan to another, we can match the activity of EM-reconstructed cells across multiple scans. This multi-scan matching has three benefits: (1) it increases the effective sample size of each cell observed in multiple scans (since we can now pool data for these cells across scans); (2) it can help detect low-SNR components (since if we find a cell in one scan we can then look for the cell in other scans — and the EM reconstruction tells us where to look); (3) we can potentially use the observed correlation structure to “fill in” estimates of the activity of neurons that are not observed on a given scan, using methods like those described in (Soudry et al., 2015).

To incorporate information across multiple scans, we first run EASE on each scan independently and compute the corresponding matching confidences (Eq. 15). Next, we select all components with high confidence levels (e.g., > 2) and create a “whitelist” of their matched EM footprints. We have high confidence that each neuron in this whitelist is well-identified and has strong signal in at least one scan; thus these neurons are likely to contribute significant signal to other scans, at the specific locations where their EM footprint intersects with the other functional imaging planes. Therefore in the next step, we run EASE again on each scan, this time only initializing EM footprints *p_i_* with IDs that appear in the whitelist. As before, these steps can be iterated.

### 4.10 Visualization of results; manual intervention

After convergence we check the results visually, using a similar strategy as in (Pnevmatikakis et al., 2016; Zhou et al., 2018). We sort the extracted components in order of their matching confidence score, defined in Eq. (15). This sorting tends to put neurons with reliable matches at the top of the list, and ambiguous neurons at the bottom of the list, allowing for a detailed checking of questionable matches. We plotted the EM footprint *p_i*_* and the functional CI footprint *a_i_* next to each other to ensure a reasonable visual match, and visualized the inferred calcium trace *c_i_* along with the projection of *a_i_* onto *Y* to look for any temporal artifacts or overly low SNR. We also found it useful to compare *a_i_* to the image generated by computing the correlation of *Y* against *c_i_*; large differences between these two images often indicate contamination by signals from a different neuron.

For any clear mismatches, we examined the top alternative matches according to the match scores. Empirically, we found that we could typically find a good match among the top-5 matches. In clear cases (e.g., when *a_i_* and the matching **p**j have large, complex spatial structures) we replaced the old mismatched *p_i*_* with the new better match. We developed a graphical user interface to facilitate this process. However, many cases remain ambiguous, particularly apical dendrites that cut through the imaging plane and only have a few pixels in *a_i_*. In these cases we might be very sure that there is an apical dendrite present at this location (i.e., the corresponding *a_i_* and *c_i_* may have large SNR), but there may be several EM components **p**j that provide plausible alternative matches, and so we simply have to declare the resulting match uncertain and pass this uncertainty on to any downstream processing stages (by keeping track of the low confidence score). Finally, in some cases no good match was found, or an oversplit was detected; in these cases we discard the non-matching or redundant component.

Next we ran a CNMF update (applying the same EM spatial constraints as before), restricted to the cells that survived the above quality checks, and then examined the residual video *Y — AC — DF — b_0_1^T^* next to the raw data *Y* and the “denoised” output AC (S1 Video). We also computed several summary images, including the correlation image (computed following (Smith and Hausser, 2010) as the mean correlation of each pixel with its neighbors) and peak-to-noise (PNR) image (i.e., the maximum minus the median signal within each pixel, divided by the noise level), of both the raw data *Y* and the residual (see Figure 3). If any clear “missing” cells were visible in the residual (either in the video or in the summary images) we ran an additional initialization step to add EM footprints *p_i_* in these spatial regions. We iterated these steps until no further discard or add steps were required.

### 4.11 Visual direction tuning

To obtain a basic characterization of the visual tuning of each recovered component, we estimated the tuning curves as the average response to moving grating stimuli (using 16 different direction angles *θ*). As in (Reimer et al., 2014) we then fit these tuning curves with a two-peak von Mises model

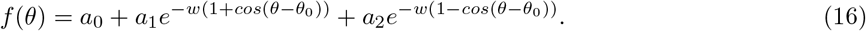

We summarize the quality of the fit with the standard R-squared value; see Figure 8 for an analysis of these “tuning curve scores.”

### 4.12 Decoding visual stimulation from the neural responses

For the decoding analysis described in section 2.4, we used a “naive Bayes” decoder (i.e., to compute the posterior distribution of the orientation *θ* we assume that each neural response is conditionally independent given *θ*), with the zero-inflated gamma (ZIG) model from Wei et al. (2019) as the encoding model (i.e., the probability model of the response of each neuron as a function of *θ*). The ZIG model is specified by three parameters: a scale parameter *α* and a shape parameter *k* for the gamma component, and the probability of non-zero responses *q*. We parameterized *α* and q as a function of orientation *θ* using neural networks (2 hidden layers, each with tanh non-linearity and 20 units). We fix the shape parameter *k* to be a constant for individual neurons (i.e., *k* is neuron-dependent but not *θ*-dependent). As in Wei et al. (2019), we optimize all the parameters by maximizing the log-likelihood using a variant of stochastic gradient descent citeadam-paper. We use a 75-20-5 train-validate-test split, and then average the test error over the 20 non-overlapping 5% splits.

### 4.13 Predicting the spatial footprint *a_i_* from the corresponding EM segment

In Figure 9 we fit a model of the form

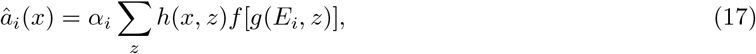

where *x* indexes 2P-resolution, two-dimensional pixels; *z* indexes EM-resolution, three-dimensional voxels; *E_i_* is the *i*-th EM component; *c_i_* is a gain factor estimated for each component *i*; *h*(*x, z*) is the (known) point-spread function (psf) operator, convolutional in lateral shifts in *x* and *z*; *g*(*E_i_*,*z*) is a featurization of the EM segment (discussed below); and *f*(*u*) is in general a nonlinear (non-negative) function of its argument *u*. We will learn *α_i_* and f from the data; *g* is fixed a priori for each model class we investigate.

We explored several featurizations. The simplest (referred to as “model 0” in Figure 9) is to simply let *f*[*g*(*E_i_, z*)] = 1 if the voxel *z* is inside the EM segment *E_i_*, and zero otherwise. In this case the predicted *a_i_* is simply the “EM footprint” *p_i_*; note that we do not estimate any parameters in f in this case.

In model 1, we let the brightness depend only on the distance from the surface of the cell, so the feature *g*(*E_i_, z*) is the distance of voxel z from the surface of EM component *E_i_*; *f*(*u*) in turn is constrained to be a unimodal function (monotonically increasing and then monotonically decreasing as a function of the scalar *u* = *g*(*E_i_*,*z*)).

In model 2, we let the brightness depend only on the distance from the center of the soma, so, as above, the feature g(E¿, z) is the distance of voxel z to the soma of EM component *E_i_*. Again, *f*(*u*) is constrained to be a unimodal function.

Finally, in model 3 we let *f*[*g*(*E_i_, z*)] be a product of models 1 and 2; i.e., the brightness of voxel z could be modulated by either the distance to the soma or the distance to the cell surface.

Each of models 1-3 were estimated by alternating constrained least-squares between the observed *a_i_* and the predicted *â_i_*. The psf is held fixed across all models. The neuron-dependent gain α_i_ is updated directly via simple regression in each iteration; updating *f* under the monotonicity constraints requires that we solve a quadratic program.

### 4.14 Validating EASE using simulated 2p video data

We tested the performance of EASE by simulating a video using the model (1). To make the simulation realistic, we took advantage of the EASE results on the real data. Specifically, the *AC* term was constructed based on the extracted neuronal signals in scan 3 and the *B* were the same as the extracted background in scan 1. The strategy of choosing results in different scans is to remove confounding correlations between AC and B in the same scan. We included 105 neurons whose temporal traces have reasonable signal quality (e.g. *skewness* ≥ 0.7). Since the extracted 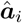 is noisy and there are some potentially some mismatches for apical dendrites, we generated each **a**i from its corresponding EM segment using the above prediction model. The resulting *a_i_* components were further normalized to match the peak values of the extracted 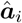. To build an empirical relationship between signal and noise, we evenly divided the ordered fluorescence values of 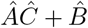 into 100 groups and stored the corresponding residuals in each group. Then we generate signal-dependent noise E at all pixels and frames by randomly drawing residuals in the corresponding groups according to the simulated *AC + B* (Figure 10A); see Buchanan et al. (2018) for a similar approach. We then applied the same pipeline as the one described in Section (2.2) to the simulated data and identified 104 components without any false positives. Given the EM segment of the missing component (this step resembles the white-list idea in the joint analysis of multiple functional imaging scans, as described in section 4.9), EASE can reliably recover its spatial footprint and temporal trace. Several quantitative and qualitative evaluations (Figure 10B-F) showed that EASE recovered the ground truth with high accuracy.

**Figure 10:**
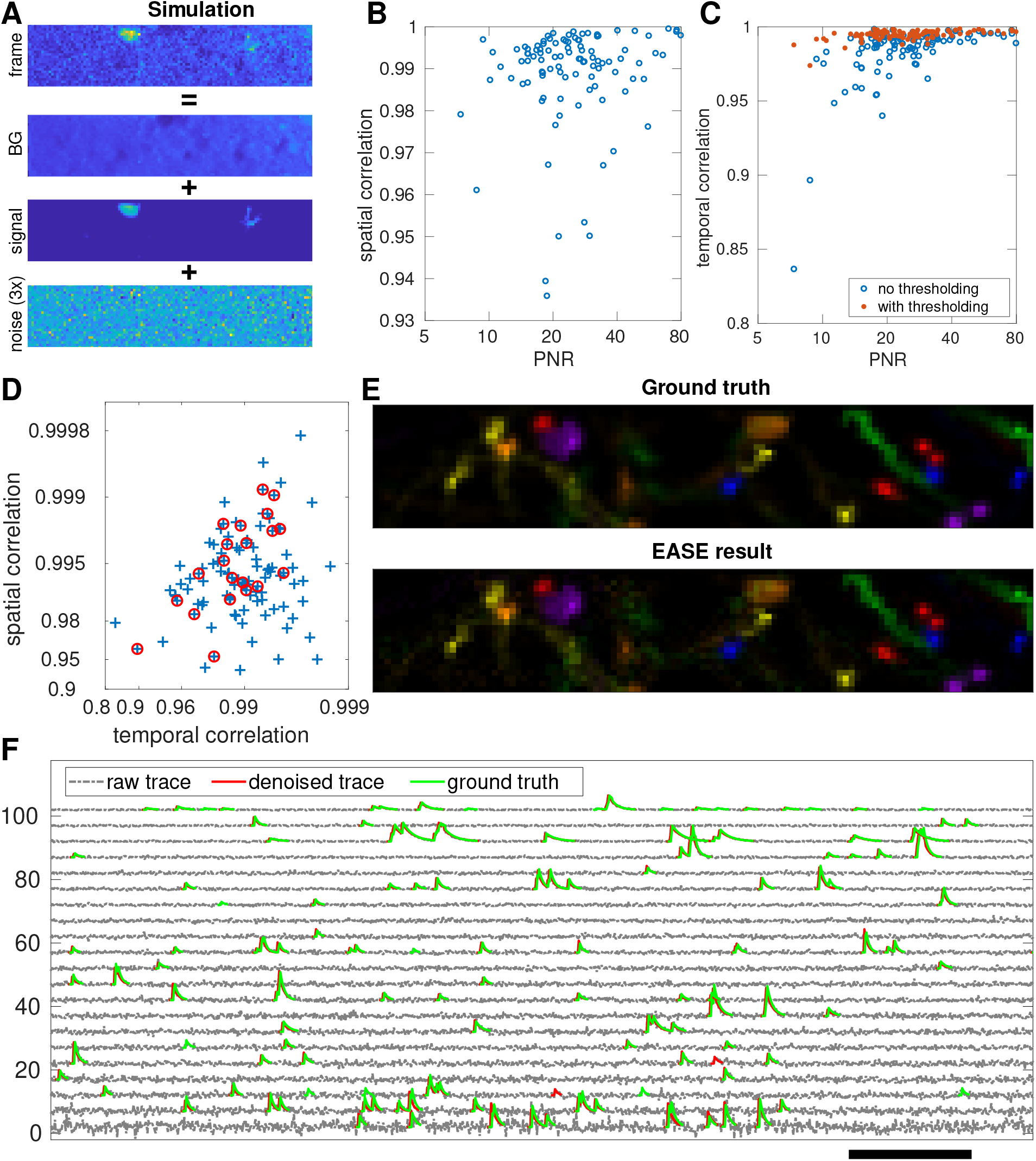
EASE correctly extracts all neurons and their temporal traces in a simulated dataset. (**A**) The process of simulating noisy videos resembling the real data; see text for full details. (**B**-**D**) The Pearson correlation coefficients between ground truth vs estimated spatial (**B**, **D**) and temporal (**C**, **D**) components. The red dots in (**C**) were computed using the thresholded traces, as a crude denoiser. The circles in (**D**) indicate example neurons in **E** and **F**. (**E**-**F**) The spatial (**E**) and the temporal (**F**) components of the selected examples. The true temporal traces were right-shifted by 100 ms and ordered by peak-to-noise ratio (PNR) for better visualization; conventions otherwise as in Figure 3B. Scale bar: 10 seconds.

### 4.15 Neural network denoising of the estimated spatial components *a_i_*

For the analysis illustrated in Figure 9, we built our spatial component denoiser using a well-established feed-forward denoising convolutional neural network (DnCNN) architecture that was developed for general natural image denoising (Zhang et al., 2017). This DnCNN takes the noisy image as an input and then outputs the noise after implicitly removing the latent clean image in the hidden layers. We first predicted the spatial components ai derived from all 453 EM segments that appeared in our EASE results using the model described in section 4.13, and then removed any images with too few nonzero pixels (here, <20 pixels), resulting in 3036 clean two-dimensional footprints among all 12 imaging planes (4 scans × 3 planes/scan). Then we randomly selected 2725 (~ 90% of the total) footprints out of the 3036 predicted footprints as training images; the other 311 were used for testing. Given these 2725 training images, we further normalized them by their maximum pixel values and augmented them by flipping, rotating and resizing (resizing factors: 2, 1.5, 1, 0.8, 0.5) the original image. Instead of using the augmented images directly, we cropped them into patches of the same size 29 × 29 and removed all-zero patches, resulting in 194581 clean image patches. The network architecture is simply a stack of *D* (*D* = 17 in this paper) convolution layers containing 64 filters of size 3 × 3 × 1. Each layer but the last utilizes rectified linear units (ReLU, max(0, o)) for the non-linearity. Batch normalization (Ioffe and Szegedy, 2015) is also added between the convolution and ReLu to speed up training (see details of the network architecture in the paper (Zhang et al., 2017)). The input to the network is the sum of a clean image patch and Gaussian white noise (σ = 0.1), and the output is expected to match the added noise, which is achieved by minimizing the mean squared error (MSE) loss function. We initialized the weights by the method in (He et al., 2015) and optimize the network with Adam (Kingma and Ba, 2014) using a mini-batch size of 256. The network can be quickly trained within 3 epochs; the learning rate is 10^-4^ after the first epoch (10^-3^).

We used PyTorch to train the network and then saved the trained network in ONNX format to be loaded by multiple deep learning platforms. We developed a unified interface to call the denoiser from either Python or Matlab. Given a noisy image, it first scales the image to match the noise level in the network training (*σ* = 0.1) and then calls the denoiser to output the noise. The difference between the input and the output is the denoised image and is scaled back to match the original noisy image.

**Table S1:** Details of processing pipeline steps for the example scan discussed in section 2.2. (*k*_1_, *k*_2_) in the column **add-neurons** indicates adding *k*_1_ neurons from the *k*_2_ EM components with the largest 1-norm ||*p_i_*||; *N* is the rank of the background model in Eq. (2).

| Iteration | add-neurons | update-variables | post-processing | # of neurons |
| --- | --- | --- | --- | --- |
| 1 | (100, 2000) | $N = 1$ | - | 100 |
| 2 | (50, 4000) | $N = 2$ | - | 150 |
| 3 | (50, 40000) | $N = 3$ | correct bad matches;<br>delete false positives | 173 |
| 4 | - | $N = 3$ | - | 173 |

## 5 Supplementary Information

### 5.1 Tables

See Table S1 for full details of the processing pipeline steps for the example scan discussed in section 2.2.

### 5.2 Videos

**S1 Video: the demixing movie of the example data in section 2.2.** All pixel values were normalized by the standard deviation of the residual. Three columns correspond to the three planes in the selected scan.

**S2 Video: the demixing movie of the 4 example neurons in Figure 5**: The spatiotemporal signal of the extracted background and all other neuronal components except the 4 selected examples were subtracted from the raw data to construct the raw signal in this video. The video was normalized in the same way as **S1 Video**. All panels use the same scaling ([-4, 4]). The yellow circle in each panel is provided as a landmark to facilitate visual comparison across panels. In the bottom panel, conventions are as in Figure 3B.

### 5.3 Experimental details

All procedures were carried out in accordance with the ethical guidelines of the National Institutes of Health and were approved by the Institutional Animal Care and Use Committee (IACUC) of Baylor College of Medicine.

#### Cranial Window

Anesthesia was induced with 3% isoflurane and maintained with 1.5% to 2% isoflurane during the surgical procedure. Mice were injected with 5-10 mg/kg ketoprofen subcutaneously at the start of the surgery. Anesthetized mice were placed in a stereotaxic head holder (Kopf Instruments) and their body temperature was maintained at 37C throughout the surgery using a homeothermic blanket system (Harvard Instruments). After shaving the scalp, bupivicane (0.05 cc, 0.5%, Marcaine) was applied subcutaneously, and after 10-20 minutes an approximately 1 cm2 area of skin was removed above the skull and the underlying fascia was scraped and removed. The wound margins were sealed with a thin layer of surgical glue (VetBond, 3M), and a 13mm stainless-steel washer clamped in the headbar was attached with dental cement (Dentsply Grip Cement). At this point, the mouse was removed from the stereotax and the skull was held stationary on a small platform by means of the newly attached headbar. Using a surgical drill and HP 1/2 burr, a 3 mm craniotomy was made centered primary visual cortex (V1; 2.7mm lateral of the midline, contacting the lambda suture), and the exposed cortex was washed with ACSF (125mM NaCl, 5mM KCl, 10mM Glucose, 10mM HEPES, 2mM CaCl2, 2mM MgSO4). The cortical window was then sealed with a 3 mm coverslip (Warner Instruments), using cyanoacrylate glue (VetBond). The mouse was allowed to recover for 1-2 hours prior to the imaging session. After imaging, the washer was released from the headbar and the mouse was returned to the home cage.

#### Widefield Imaging

Prior to two-photon imaging, we acquired a low-magnification image of the 3mm craniotomy under standard illumination. The location of the subsequent two-photon field of view could then be identified in this image based on surface vasculature. The location of the target two-photon imaging site in V1 was determined by retinotopic mapping using intrinsic signal imaging or GCaMP6 imaging at low magnification.

#### Two-photon Imaging

Two-photon imaging was performed in V1, in a 400 x 400 x 200 um volume with the superficial surface of the volume at the border of L1 and L2/3, approximately 100 um below the pia. Imaging data was collected with a resonant scanning microscope (ThorLabs) and software (Scanimage 5.1, Vidrio). Nine scans were collected in total, starting superficially and moving deeper into the cortex with each subsequent scan. During each 30-minute scan, a piezo controlled manipulator (PI-726, Physik Instruments) moved the microscope objective between three different z-planes (“slices”). These three slices were separated by an average of 8 um by the piezo, and each slice was imaged at 14.8313 frames per second. (We refer to each trio of sequential slices as a one imaging “scan”.) Since we collected 9 scans with 3 slices/scan, we had 27 slices in total to span the 200um depth of the overall 400 x 400 x 200 um imaging volume. Two color channels were recorded: Channel one was GCaMP6 calcium imaging and channel two was blood vessels labeled with red dye (Sulfarhodamine 101). Thus each functional scan is a 256 x 256 x 2 channels x 3 slices x 27300 volume 16-bit TIFF stack.

The mouse was head-restrained but could walk on a treadmill during imaging. While we were imaging, we collected treadmill speed at 200Hz (recorded in an HDF5 file) and we recorded a movie of the mouse’s eye at 640 x 480 @20 Hz (recorded as uncompressed .AVI). Visual stimuli were presented at 60 fps, and synchronized to imaging and behavioral data via a photodiode which recorded the timing of each stimulus frame. For each 30 minute and 40 second scan (27300 volumes at 14.8313 volumes per second) we presented 30 one-minute trials of a colored-noise stimulus (Niell and Stryker, 2008) interspersed with periods of coherent motion of oriented noise. Each one minute trial contained 16 stationary-moving-stationary blocks, with a different direction presented in each block, pseudorandomly-ordered.

To facilitate alignment with EM, at the beginning of the experiment we collected a high-resolution structural stack of the imaging volume with the same field of view and xy location as the functional scans. This stack began 310 um deep and ended at the cortical surface, in one micron steps. This stack is saved as a 512×512×2 channels x 310 slices 16-bit TIFF stack.

### 5.4 Algorithm for solving (P-S)(P-T)(P-B)

There are already existing algorithms for solving the optimization problems that are the same as or similar to the subproblems (P-S)(P-T)(P-B). We summarize the algorithms used in this work in Algorithm (3) and briefly describe them below.

The same problem of (P-S) has been discussed in (Friedrich et al., 2017b) and (Zhou et al., 2018). These two papers generated the spatial support of each *a_i_* based on its previous estimation, while here we use *supp*(***p**_i*_*) directly. The algorithm used there is modified from fastHALS (Cichocki and Phan, 2009). Note that the estimation of *A* is independent of 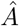 because A only accounts for the fluctuating signals. We estimate A first and then update ***b***_0_ using the closed-form expression 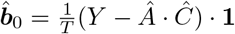.

(Zhou et al., 2018) has a similar (P-T) problem as here except the weighting term 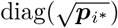, thus we can reuse the algorithm by trivially including the weighting term. We iteratively update all neurons, and for each neuron we first compute its unconstrained estimate

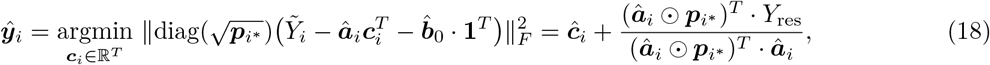

followed by deconvolving and denoising 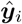 to infer the denoised trace 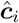 and the deconvolved signal 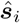 (Friedrich et al., 2017b). Once 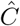 is estimated, we also update *b*_0_ as 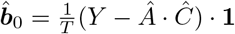.

The constraint *F* · 1 = 0 in (P-B) automatically yields a closed-form estimate of *b*_0_ as 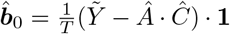. Then (P-B) becomes a standard singular value decomposition (SVD) problem and the solution corresponds to the top-*N* singular components of 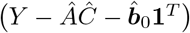.

**Algorithm 3.**
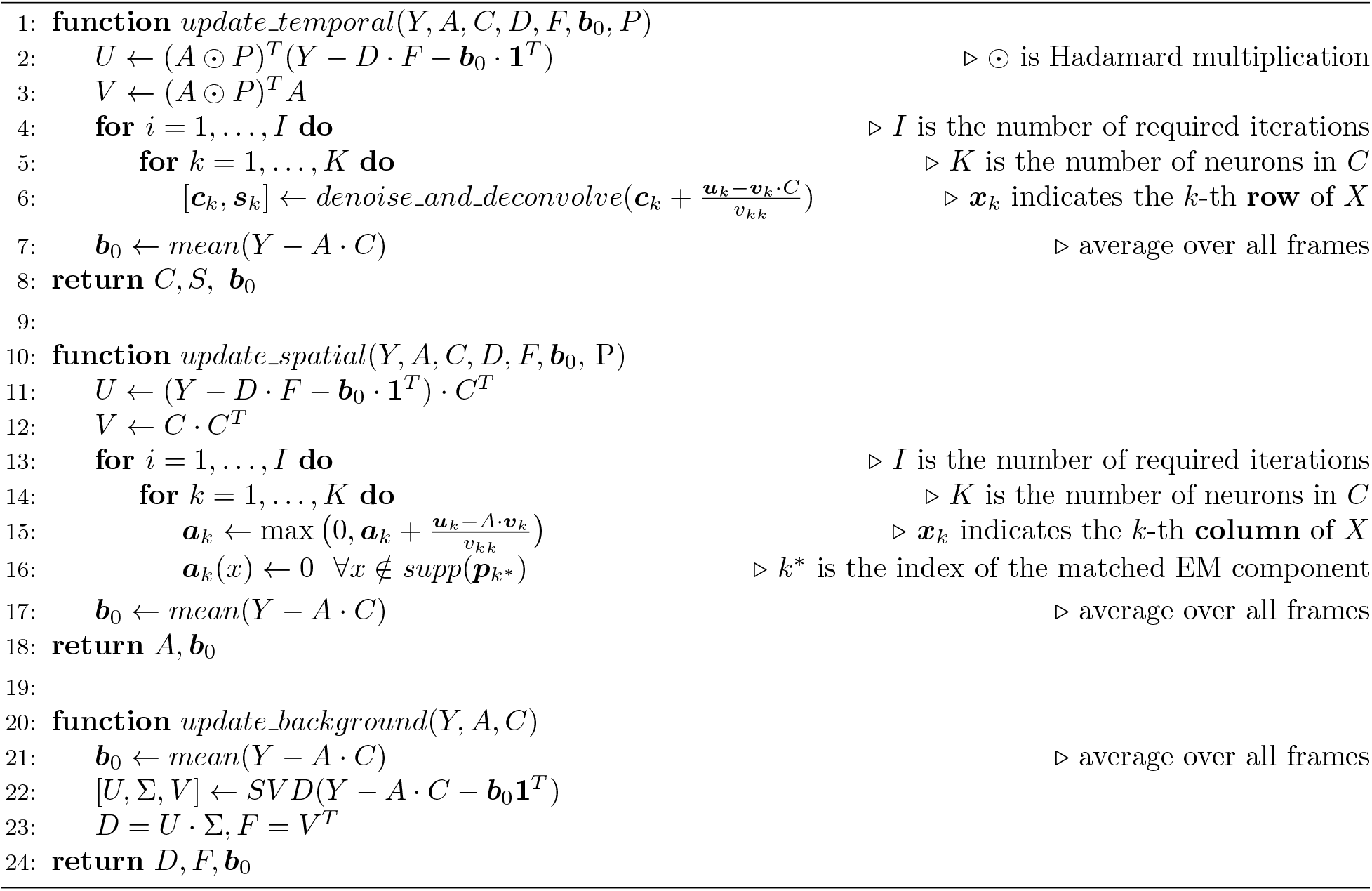
Functions for updating EASE variables

### 5.5 Derivation of Eq. (8)

Each EM segment ***p**_i*_* divides the FOV into two areas: pixels within and pixels outside of *supp*(***p**_i*_*). By ignoring the noise term *E* and centering all temporal signals in our matrix factorization model (Eq. 1), we can rewrite the model in these two areas separately:

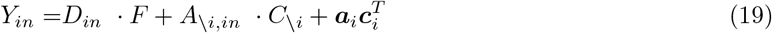

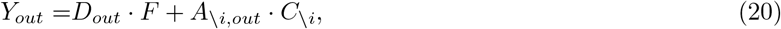

where ***a**_i_* and ***c**_i_* are the spatial and the temporal components for the neuron corresponding to ***p**_i*_. Y_out_* in Eq. (20) has no signals from ***c**_i_* because *supp*(***a**_i_*) ⊆ *supp*(***p**_i*_*).

If we apply SVD to decompose *Y_out_* as *U · S · V^T^*, then *F* and *C_\i_* can largely be represented in the *V^T^* basis. Then the contamination term *D_in_ · F* + *A_/i,in_* · *C_\i_* can be simplified as *W · V^T^* for some weight matrix *W*, yielding Eq. (8) by substituing it into Eq. (19). In practice, we only choose pixels surrounding *supp*(*p_i_*) to construct *Y_out_* and use a truncated version of V because they are enough to approximate the contaminating signals in *Y_in_.*

### 5.6 Initialization of single neural components

In section 4.7.2 we defined an approach for initializing functional components from the EM footprints *p_i_*, along with a simpler baseline semi-NMF method. The main difference between these two approaches is that the EASE initialization regresses out the contribution of cells that overlap with ***p**_i_* but display significant signal outside the support of ***p**_i_*. We also minimize a ***p**_i_*-weighted RSS (wRSS, Eq. (11)), instead of the standard RSS, to further reduce nearby contaminations. We validated the necessity of these two novel contributions by simulation, where we added one neuron’s spatiotemporal signal to the calcium imaging data of one scan and tried to recover this neuron’s spatial and temporal components with different initialization algorithms. To make the simulation realistic and avoid confounding correlations, we generated the added neural signal from a neuron extracted from a different scan taken at a different time. Figure S1AB shows the comparison of different algorithms; the proposed algorithm for EASE performs the best visually. Next we varied the SNR level of the added neural signal by multiplying its temporal component by a scale factor, and quantified performance by computing the correlations between the ground-truth and the initialized results (Figure S1CD). Again, the proposed EASE initialization outperformed the alternative approaches consistently.

**Figure S1:**
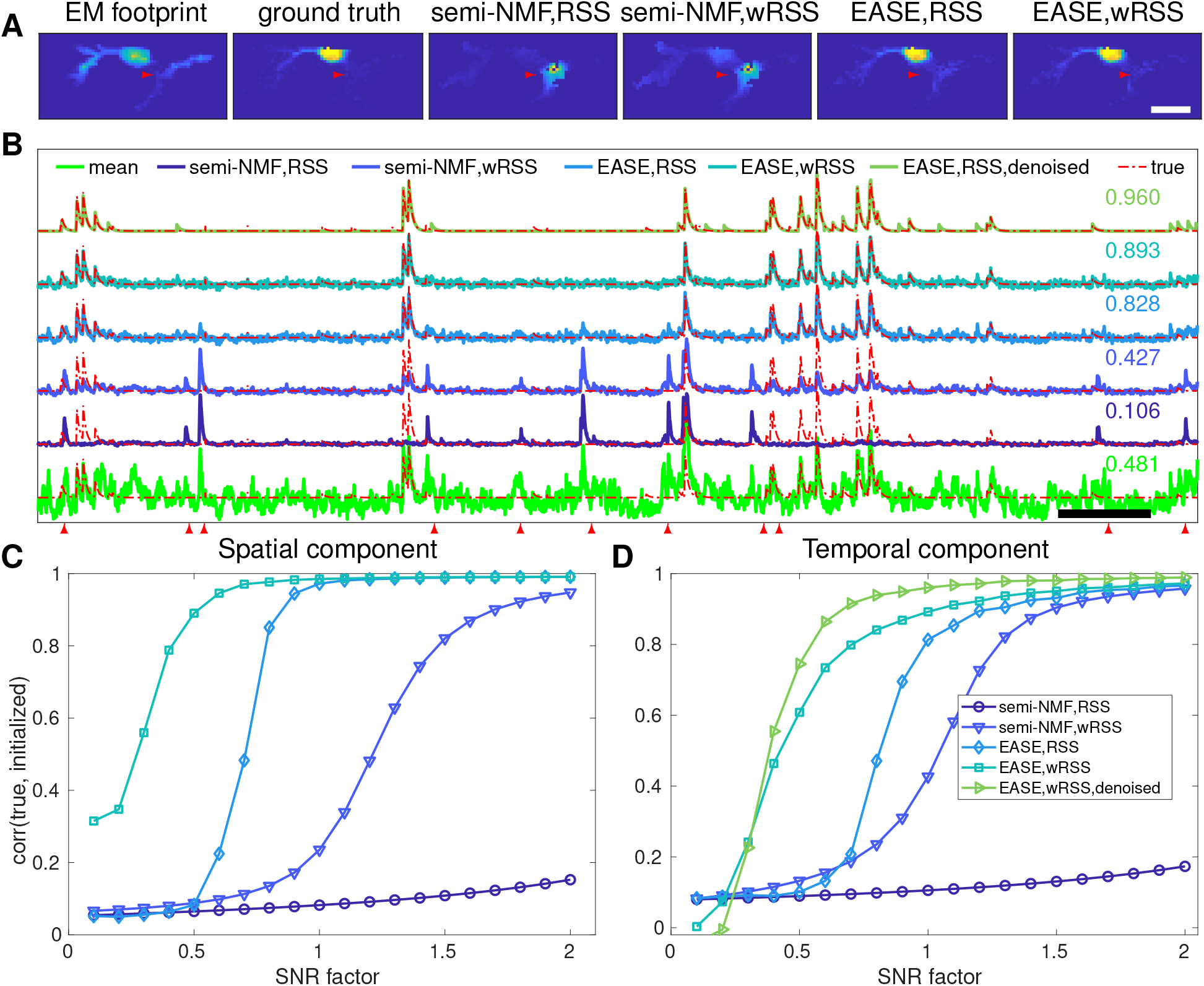
Comparison of different initialization algorithms. (**A**) The ground truth and the initialized spatial footprints of the selected neuron. For simplicity, only one of the three imaging planes shown here (scale bar: 20 um). (**B**) The initialized temporal traces and their correlations with the ground-truth. (Scale bar: 20 seconds). Red arrows in (**A**) and (**B**) highlight the locations or bins with large discrepancies between the estimate and ground truth. (**C** and **D**) The correlation between the ground-truth and the inferred components under different SNR levels. EASE using weighted RSS (instead of unweighted RSS) and applying denoising to the temporal trace consistently achieves the best performance in this simulation; similar results are seen when adding other ground truth components (data not shown)

## Footnotes

1 https://www.mathworks.com/matlabcentral/fileexchange/24086-polygon2voxel

2 To derive Eq. (11) from Eq. (10), we use the following: 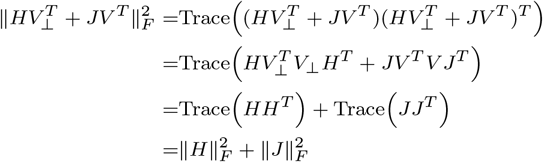

## Notes

https://github.com/zhoupc/sharing/blob/master/data/EASE.md

## References

Alon, S., Huynh, G. H., and Boyden, E. S. (2018). Expansion microscopy: Enabling single cell analysis in intact biological systems. The FEBS journal.

Bae, J. A., Mu, S., Kim, J. S., Turner, N. L., Tartavull, I., Kemnitz, N., Jordan, C. S., Norton, A. D., Silversmith, W. M., Prentki, R., Sorek, M., David, C., Jones, D. L., Bland, D., Sterling, A. L. R., Park, J., Briggman, K. L., and Seung, H. S. (2018). Digital Museum of Retinal Ganglion Cells with Dense Anatomy and Physiology. Cell, 173(5):1293–1306.e19.

Begemann, I. and Galic, M. (2016). Correlative Light Electron Microscopy: Connecting Synaptic Structure and Function. Frontiers in Synaptic Neuroscience, 8.

Blazquez-Llorca, L., Hummel, E., Zimmerman, H., Zou, C., Burgold, S., Rietdorf, J., and Herms, J. (2015). Correlation of two-photon in vivo imaging and FIB/SEM microscopy. Journal of Microscopy, 259(2):129–136.

Bock, D. D., Lee, W.-c. A., Kerlin, A. M., Andermann, M. L., Hood, G., Wetzel, A. W., Yurgenson, S., Soucy, E. R., Kim, H. S., and Reid, R. C. (2011). Network anatomy and in vivo physiology of visual cortical neurons. Nature, 471(7337):177–182.

Boyd, S. and Vandenberghe, L. (2004). Convex Optimization. Cambridge university press.

Briggman, K. L., Helmstaedter, M., and Denk, W. (2011). Wiring specificity in the direction-selectivity circuit of the retina. Nature, 471(7337):183–8.

Buchanan, E. K., Kinsella, I., Zhou, D., Zhu, R., Zhou, P., Gerhard, F., Ferrante, J., Ma, Y., Kim, S., Shaik, M., Liang, Y., Lu, R., Reimer, J., Fahey, P., Muhammad, T., Dempsey, G., Hillman, E., Ji, N., Toias, A., and Paninski, L. (2018). Penalized matrix decomposition for denoising, compression, and improved demixing of functional imaging data. bioRxiv, page 334706.

Charles, A. S., Song, A., Gauthier, J. L., Pillow, J. W., and Tank, D. W. (2019). Neural Anatomy and Optical Microscopy (NAOMi) Simulation for evaluating calcium imaging methods. bioRxiv, page 726174.

Cichocki, A. and Phan, A. H. (2009). Fast local algorithms for large scale nonnegative matrix and tensor factorizations. IEICE Transactions on Fundamentals of Electronics, Communications and Computer Sciences, E92-A(3):708–721.

Cichocki, A., Zdunek, R., and Amari, S.-i. (2007). Hierarchical ALS algorithms for nonnegative matrix and 3D tensor factorization. In International Conference on Independent Component Analysis and Signal Separation, pages 169–176. Springer.

de Boer, P., Hoogenboom, J. P., and Giepmans, B. N. G. (2015). Correlated light and electron microscopy: Ultrastructure lights up! Nature Methods, 12(6):503–513.

Ding, H., Smith, R. G., Poleg-Polsky, A., Diamond, J. S., and Briggman, K. L. (2016). Species-specific wiring for direction selectivity in the mammalian retina. Nature, 535(7610):105–110.

Donoho, D. (2017). 50 Years of Data Science. Journal of Computational and Graphical Statistics, 26(4):745–766.

Dorkenwald, S., Turner, N. L., Macrina, T., Lee, K., Lu, R., Wu, J., Bodor, A. L., Bleckert, A. A., Brittain, D., Kemnitz, N., Silversmith, W. M., Ih, D., Zung, J., Zlateski, A., Tartavull, I., Yu, S.-C., Popovych, S., Wong, W., Castro, M., Jordan, C. S., Wilson, A. M., Froudarakis, E., Buchanan, J., Takeno, M., Torres, R., Mahalingam, G., Collman, F., Schneider-Mizell, C., Bumbarger, D. J., Li, Y., Becker, L., Suckow, S., Reimer, J., Tolias, A. S., da Costa, N. M., Reid, R. C., and Seung, H. S. (2019). Binary and analog variation of synapses between cortical pyramidal neurons. bioRxiv.

Drawitsch, F., Karimi, A., Boergens, K. M., and Helmstaedter, M. (2018). FluoEM, virtual labeling of axons in three-dimensional electron microscopy data for long-range connectomics. eLife, 7:e38976.

Friedrich, J., Yang, W., Soudry, D., Mu, Y., Ahrens, M. B., Yuste, R., Peterka, D. S., and Paninski, L. (2017a). Multi-scale approaches for high-speed imaging and analysis of large neural populations. PLoS computational biology, 13(8):e1005685.

Friedrich, J., Zhou, P., and Paninski, L. (2017b). Fast online deconvolution of calcium imaging data. PLOS Computational Biology, 13(3):e1005423.

Gao, R., Asano, S. M., Upadhyayula, S., Pisarev, I., Milkie, D. E., Liu, T.-L., Singh, V., Graves, A., Huynh, G. H., Zhao, Y., Bogovic, J., Colonell, J., Ott, C. M., Zugates, C., Tappan, S., Rodriguez, A., Mosaliganti, K. R., Sheu, S.-H., Pasolli, H. A., Pang, S., Xu, C. S., Megason, S. G., Hess, H., Lippincott-Schwartz, J., Hantman, A., Rubin, G. M., Kirchhausen, T., Saalfeld, S., Aso, Y., Boyden, E. S., and Betzig, E. (2019). Cortical column and whole-brain imaging with molecular contrast and nanoscale resolution. Science, 363(6424):eaau8302.

Giovannucci, A., Friedrich, J., Gunn, P., Kalfon, J., Brown, B. L., Koay, S. A., Taxidis, J., Najafi, F., Gauthier, J. L., Zhou, P., Khakh, B. S., Tank, D. W., Chklovskii, D. B., and Pnevmatikakis, E. A. (2019). CaImAn an open source tool for scalable calcium imaging data analysis. eLife, 8.

Hayworth, K. J., Xu, C. S., Lu, Z., Knott, G. W., Fetter, R. D., Tapia, J. C., Lichtman, J. W., and Hess, H. F. (2015). Ultrastructurally smooth thick partitioning and volume stitching for large-scale connectomics. Nature Methods, 12(4):319–322.

He, K., Zhang, X., Ren, S., and Sun, J. (2015). Delving Deep into Rectifiers: Surpassing Human-Level Performance on ImageNet Classification. In 2015 IEEE International Conference on Computer Vision (ICCV), pages 1026–1034, Santiago, Chile. IEEE.

Helmstaedter, M., Briggman, K. L., Turaga, S. C., Jain, V., Seung, H. S., and Denk, W. (2013). Connectomic reconstruction of the inner plexiform layer in the mouse retina. Nature, 500(7461):168–174.

Hildebrand, D. G. C., Cicconet, M., Torres, R. M., Choi, W., Quan, T. M., Moon, J., Wetzel, A. W., Scott Champion, A., Graham, B. J., Randlett, O., Plummer, G. S., Portugues, R., Bianco, I. H., Saalfeld, S., Baden, A. D., Lillaney, K., Burns, R., Vogelstein, J. T., Schier, A. F., Lee, W. C. A., Jeong, W. K., Lichtman, J. W., and Engert, F. (2017). Whole-brain serial-section electron microscopy in larval zebrafish. Nature, 545(7654):345–349.

Hoffman, D. P., Shtengel, G., Xu, C. S., Campbell, K. R., Freeman, M., Wang, L., Milkie, D. E., Pasolli, H. A., Iyer, N., Bogovic, J. A., et al. (2020). Correlative three-dimensional super-resolution and block-face electron microscopy of whole vitreously frozen cells. Science, 367(6475).

Ioffe, S. and Szegedy, C. (2015). Batch Normalization: Accelerating Deep Network Training by Reducing Internal Covariate Shift. arXiv:1502.03167 [cs].

Januszewski, M., Kornfeld, J., Li, P. H., Pope, A., Blakely, T., Lindsey, L., Maitin-Shepard, J., Tyka, M., Denk, W., and Jain, V. (2018). High-precision automated reconstruction of neurons with flood-filling networks. Nature Methods, 15(8):605–610.

Kasthuri, N., Hayworth, K. J., Berger, D. R., Schalek, R. L., Conchello, J. A., Knowles-Barley, S., Lee, D., Vazquez-Reina, A., Kaynig, V., and Jones, T. R. (2015). Saturated reconstruction of a volume of neocortex. Cell, 162(3):648–661.

Kim, J. S., Greene, M. J., Zlateski, A., Lee, K., Richardson, M., Turaga, S. C., Purcaro, M., Balkam, M., Robinson, A., Behabadi, B. F., Campos, M., Denk, W., Seung, H. S., and the EyeWirers (2014). Space-time wiring specificity supports direction selectivity in the retina. Nature, 509(7500):331–336.

Kingma, D. P. and Ba, J. (2014). Adam: A Method for Stochastic Optimization. arXiv:1412.6980 [cs].

Lee, W. C. A., Bonin, V., Reed, M., Graham, B. J., Hood, G., Glattfelder, K., and Reid, R. C. (2016). Anatomy and function of an excitatory network in the visual cortex. Nature, 532(7599):370–374.

Lees, R. M., Peddie, C. J., Collinson, L. M., Ashby, M. C., and Verkade, P. (2017). Chapter 12 - Correlative two-photon and serial block face scanning electron microscopy in neuronal tissue using 3D near-infrared branding maps. In Müller-Reichert, T. and Verkade, P., editors, Methods in Cell Biology, volume 140 of Correlative Light and Electron Microscopy III, pages 245–276. Academic Press.

Maco, B., Cantoni, M., Holtmaat, A., Kreshuk, A., Hamprecht, F. A., and Knott, G. W. (2014a). Semiautomated correlative 3D electron microscopy of in vivo-imaged axons and dendrites. Nature Protocols, 9(6):1354–1366.

Maco, B., Holtmaat, A., Cantoni, M., Kreshuk, A., Straehle, C. N., Hamprecht, F. A., and Knott, G. W. (2013). Correlative In Vivo 2 Photon and Focused Ion Beam Scanning Electron Microscopy of Cortical Neurons. PLoS ONE, 8(2):e57405.

Maco, B., Holtmaat, A., Jorstad, A., Fua, P., and Knott, G. W. (2014b). Chapter 16 - Correlative In Vivo 2-Photon Imaging and Focused Ion Beam Scanning Electron Microscopy: 3D Analysis of Neuronal Ultrastructure. In Müller-Reichert, T. and Verkade, P., editors, Methods in Cell Biology, volume 124 of Correlative Light and Electron Microscopy II, pages 339–361. Academic Press.

Madisen, L., Garner, A. R., Shimaoka, D., Chuong, A. S., Klapoetke, N. C., Li, L., van der Bourg, A., Niino, Y., Egolf, L., Monetti, C., Gu, H., Mills, M., Cheng, A., Tasic, B., Nguyen, T. N., Sunkin, S. M., Benucci, A., Nagy, A., Miyawaki, A., Helmchen, F., Empson, R. M., Knopfel, T., Boyden, E. S., Reid, R. C., Carandini, M., and Zeng, H. (2015). Transgenic mice for intersectional targeting of neural sensors and effectors with high specificity and performance. Neuron, 85(5):942–958.

Morgan, J. L., Berger, D. R., Wetzel, A. W., and Lichtman, J. W. (2016). The Fuzzy Logic of Network Connectivity in Mouse Visual Thalamus. Cell, 165(1):192–206.

Motta, A., Berning, M., Boergens, K. M., Staffler, B., Beining, M., Loomba, S., Hennig, P., Wissler, H., and Helmstaedter, M. (2019). Dense connectomic reconstruction in layer 4 of the somatosensory cortex. Science.

Mukamel, E. A., Nimmerjahn, A., and Schnitzer, M. J. (2009). Automated Analysis of Cellular Signals from Large-Scale Calcium Imaging Data. Neuron, 63(6):747–760.

Niell, C. M. and Stryker, M. P. (2008). Highly Selective Receptive Fields in Mouse Visual Cortex. Journal of Neuroscience, 28(30):7520–7536.

Pachitariu, M., Stringer, C., Schrüder, S., Dipoppa, M., Rossi, L. F., Carandini, M., and Harris, K. D. (2016). Suite2p: Beyond 10,000 neurons with standard two-photon microscopy. bioRxiv, pages 061507–061507.

Paninski, L. and Cunningham, J. P. (2018). Neural data science: Accelerating the experiment-analysis-theory cycle in large-scale neuroscience. Current Opinion in Neurobiology, 50:232–241.

Petersen, A., Simon, N., and Witten, D. (2018). SCALPEL: Extracting neurons from calcium imaging data. The Annals of Applied Statistics, 12(4):2430–2456.

Pnevmatikakis, E. A., Soudry, D., Gao, Y., Machado, T. A., Merel, J., Pfau, D., Reardon, T., Mu, Y., Lacefield, C., Yang, W., Ahrens, M., Bruno, R., Jessell, T. M., Peterka, D. S., Yuste, R., and Paninski, L. (2016). Simultaneous denoising, deconvolution, and demixing of calcium imaging data. Neuron, 89(2):285–299.

Reimer, J., Froudarakis, E., Cadwell, C. R., Yatsenko, D., Denfield, G. H., and Tolias, A. S. (2014). Pupil Fluctuations Track Fast Switching of Cortical States during Quiet Wakefulness. Neuron, 84(2):355–362.

Smith, S. L. and Hausser, M. (2010). Parallel processing of visual space by neighboring neurons in mouse visual cortex. Nature Neuroscience, 13(9):1144–1149.

Soltanian-Zadeh, S., Sahingur, K., Blau, S., Gong, Y., and Farsiu, S. (2019). Fast and robust active neuron segmentation in two-photon calcium imaging using spatiotemporal deep learning. Proceedings of the National Academy of Sciences, 116(17):8554–8563.

Soudry, D., Keshri, S., Stinson, P., Oh, M.-h., Iyengar, G., and Paninski, L. (2015). Efficient” shotgun” inference of neural connectivity from highly sub-sampled activity data. PLoS computational biology, 11(10):e1004464.

Takemura, S.-y., Nern, A., Chklovskii, D. B., Scheffer, L. K., Rubin, G. M., and Meinertzhagen, I. A. (2017). The comprehensive connectome of a neural substrate for ‘ON’ motion detection in Drosophila. eLife, 6.

Vishwanathan, A., Daie, K., Ramirez, A. D., Lichtman, J. W., Aksay, E. R., and Seung, H. S. (2017). Electron microscopic reconstruction of functionally identified cells in a neural integrator. Current Biology, 27(14):2137–2147.

Vogelstein, J. T., Packer, A. M., Machado, T. A., Sippy, T., Babadi, B., Yuste, R., and Paninski, L. (2010). Fast non-negative deconvolution for spike train inference from population calcium imaging. Journal of Neurophysiology, 104(6):3691–3704.

Wei, X.-X., Zhou, D., Grosmark, A., Ajabi, Z., Sparks, F., Zhou, P., Brandon, M., Losonczy, A., and Paninski, L. (2019). A zero-inflated gamma model for post-deconvolved calcium imaging traces. bioRxiv, page 637–652.

Zhang, K., Zuo, W., Chen, Y., Meng, D., and Zhang, L. (2017). Beyond a Gaussian Denoiser: Residual Learning of Deep CNN for Image Denoising. IEEE Transactions on Image Processing, 26(7):3142–3155.

Zheng, Z., Lauritzen, J. S., Perlman, E., Robinson, C. G., Nichols, M., Milkie, D., Torrens, O., Price, J., Fisher, C. B., Sharifi, N., Calle-Schuler, S. A., Kmecova, L., Ali, I. J., Karsh, B., Trautman, E. T., Bogovic, J., Hanslovsky, P., Jefferis, G. S. X. E., Kazhdan, M., Khairy, K., Saalfeld, S., Fetter, R. D., and Bock, D. D. (2017). A Complete Electron Microscopy Volume Of The Brain Of Adult Drosophila melanogaster. bioRxiv.

Zhou, P., Resendez, S. L., Rodriguez-Romaguera, J., Jimenez, J. C., Neufeld, S. Q., Giovannucci, A., Friedrich, J., Pnevmatikakis, E. A., Stuber, G. D., Hen, R., Kheirbek, M. A., Sabatini, B. L., Kass, R. E., and Paninski, L. (2018). Efficient and accurate extraction of in vivo calcium signals from microendoscopic video data. eLife, 7:1–14.

